# SKELETAL MyBP-C ISOFORMS TUNE THE MOLECULAR CONTRACTILITY OF DIVERGENT SKELETAL MUSCLE SYSTEMS

**DOI:** 10.1101/676601

**Authors:** Amy Li, Shane Nelson, Sheema Rahmanseresht, Filip Braet, Anabelle S. Cornachione, Samantha Beck Previs, Thomas S. O’Leary, James W. McNamara, Dilson E. Rassier, Sakthivel Sadayappan, Michael J. Previs, David M. Warshaw

**Affiliations:** Department of Molecular Physiology and Biophysics, Cardiovascular Research Institute, University of Vermont, Burlington, VT 05405 USA; Discipline of Anatomy & Histology, School of Medical Sciences, The University of Sydney, Sydney, NSW 2006 Australia; Australian Centre for Microscopy & Microanalysis, The University of Sydney, Sydney, NSW 2006 Australia; Department of Physiological Science, Federal University of São Carlos, São Carlos, Brazil; Department of Kinesiology and Physical Education, McGill University, Montreal, Canada; Heart, Lung and Vascular Institute, Department of Internal Medicine, Division of Cardiovascular Health and Disease, University of Cincinnati, Cincinnati, OH 45267, USA

**Keywords:** muscle contraction, myosin thick filament, actin thin filament, mass spectrometry, *in vitro* motility, Monte Carlo simulation, soleus, extensor digitorum longus, calcium regulation

## Abstract

Skeletal muscle myosin-binding protein C (MyBP-C) is a myosin thick filament-associated protein; localized through its C terminus to distinct regions (C-zones) of the sarcomere. MyBP-C modulates muscle contractility, presumably through its N terminus extending from the thick filament and interacting with either the myosin head region and/or the actin thin filament. Two isoforms of MyBP-C (fast- and slow-type) are expressed in a muscle-type specific manner. Are the expression, localization, and Ca^2+^-dependent modulatory capacities of these isoforms different in fast-twitch extensor digitorum longus (EDL) and slow-twitch soleus (SOL) muscles derived from Sprague-Dawley rats? By mass spectrometry, four MyBP-C isoforms (one fast-type MyBP-C and three N-terminally spliced slow-type MyBP-C) were expressed in EDL but only the three slow-type MyBP-C isoforms in SOL. Using EDL and SOL native thick filaments in which the MyBP-C stoichiometry and localization are preserved, native thin filament sliding over these thick filaments showed that only in the C-zone, MyBP-C Ca^2+^-sensitizes the thin filament and slows thin filament velocity. These modulatory properties depended on MyBP-C’s N-terminus, as N-terminal proteolysis attenuated MyBP-C’s functional capacities. To determine each MyBP-C isoform’s contribution to thin filament Ca^2+^-sensitization and slowing in the C-zone, we used a combination of *in vitro* motility assays using expressed recombinant N-terminal fragments and *in silico* mechanistic modeling. Our results suggest that each skeletal MyBP-C isoform’s N terminus is functionally distinct and has modulatory capacities that depend on the muscle-type in which they are expressed, providing the potential for molecular tuning of skeletal muscle performance through differential MyBP-C expression.

**SIGNIFICANCE:** Myosin-binding protein C (MyBP-C) is a critical component of the skeletal muscle sarcomere, muscle’s smallest contractile unit. MyBP-C’s importance is evident by genetic mutations leading to human myopathies, such as distal arthrogryposis (i.e. club foot). However, the molecular basis of MyBP-C’s functional impact on skeletal muscle contractility is far from certain. Complicating matters further is the expression of fast- and slow-type MyBP-C isoforms that depend on whether the muscle is fast- or slow-twitch. Using multi-scale proteomic, biophysical and mathematical modeling approaches, we define the expression, localization, and modulatory capacities of these distinct skeletal MyBP-C isoforms in rat skeletal muscles. Each MyBP-C isoform appears to modulate muscle contractility differentially; providing the capacity to fine-tune muscle’s mechanical performance as physiological demands arise.

## INTRODUCTION

Skeletal muscle myosin-binding protein C (MyBP-C) is a ~130 kDa, thick filament-associated protein and a key modulator of muscle contractility (1). Its importance is emphasized by mutations in its genes resulting in human skeletal muscle myopathies, such as distal arthrogryposis (2, 3). MyBP-C has a modular structure, consisting of 7 immunoglobulin (Ig) and 3 fibronectin-like (Fn) domains, C1 to C10 (Fig. 1B). Through its C-terminal domain interactions with the thick filament backbone (4), MyBP-C is reportedly localized to 7-9 stripes within a distinct central region (i.e. the C-zone) of the sarcomere (Fig. 1A), muscle’s smallest contractile unit (5). At its other end, the N terminus extends away from the thick filament, allowing it to interact transiently with either the myosin head region (6–9) and/or actin thin filament (10–14).

**Figure 1.**
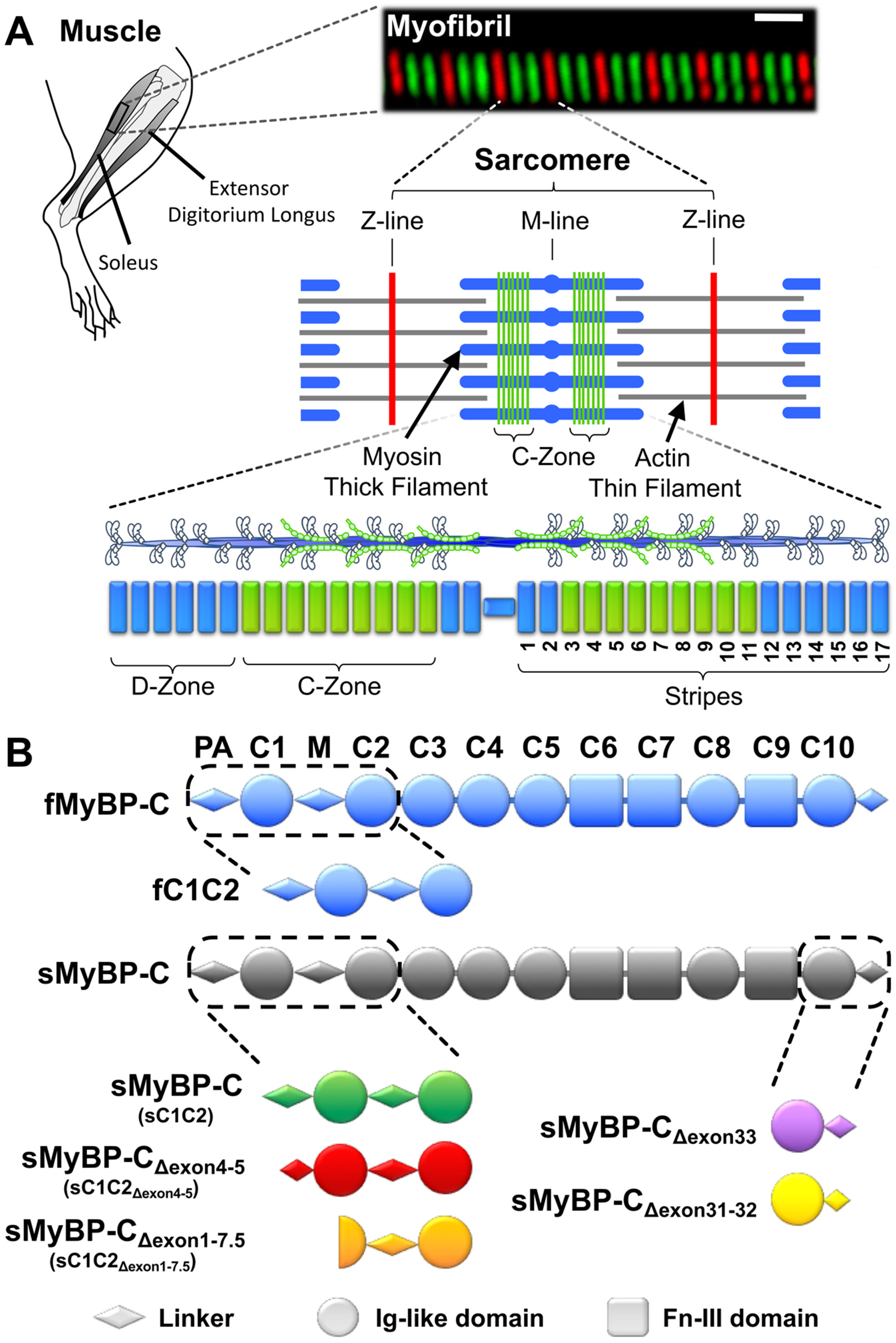
MyBP-C isoforms and their localization in rat skeletal muscle. **(A)** In this study contractile protein preparations were obtained from the rat hind limb soleus and extensor digitorum longus. An immuno-fluorescent image of a soleus myofibril composed of 8 sarcomeres, whose boundaries are identified by antibodies to the Z-line protein, α-actinin (red). Antibodies against slow-type MyBP-C (green) highlight the distinct regions (C-Zones) in the sarcomere center where MyBP-C is localized. The sarcomere is comprised of myosin thick filaments (blue) and actin thin filaments (grey). Striated muscle myosin forms bipolar thick filaments where each half thick filament consists of 17 stripes that demarcate the 43 nm myosin head helical repeat. MyBP-C within the C-Zone occupies ~9 stripes near the thick filament center, where the ends of the thick filament are devoid of MyBP-C (D-Zone). **(B)** Skeletal MyBP-C consists of seven immunoglobulin-like (Ig-like, circles) and three fibronectin-III (Fn-III, rectangles) domains named C1 through C10. A proline-alanine rich domain (PA) precedes the C1 domain and an M-domain linker (M) exists between the C1 and C2 domains. In skeletal muscle, one fast-type MyBP-C and multiple slow-type MyBP-C splice isoforms are present. Slow-type MyBP-C is alternatively spliced at its N- and C-terminus. See Figs. S1 and S2 for isoform specific sequences.

The functional impact of MyBP-C was first demonstrated when partial extraction of the whole molecule in skinned skeletal muscle fibers resulted in both increased force production and shortening velocity at submaximal calcium levels (15, 16). Based on more recent *in vitro* data using expressed MyBP-C fragments, MyBP-C’s N-terminal domain interactions with the actin thin filament and/or myosin head may be the primary mechanistic modulators of muscle contractility (17, 18). Specifically, N-terminal fragment binding to the thin filament can sensitize the thin filament to calcium by shifting the position of tropomyosin out of the “blocked” state to facilitate myosin binding to actin (18). In addition, MyBP-C can act as a “brake” to reduce sarcomere shortening (19) and thin filament sliding velocity (17, 18) through its interaction with either the thin filament and/or myosin head region. Although extremely informative, the *in vitro* N-terminal fragment studies were performed with contractile proteins isolated from heterologous species or muscle tissue, creating non-native simplified contractile model systems. To complicate matters further, two mammalian skeletal muscle MyBP-C isoforms exist that are encoded by separate genes, *MYBPC1* and *MYBPC2*. These isoforms were historically named slow-type skeletal MyBP-C (*MYBPC1*) and fast-type skeletal MyBP-C (*MYBPC2*) based on the twitch speed of the muscle in which they were expressed (20). Recent studies suggest that the number of MyBP-C isoforms and their muscle type expression profiles are far more complex due to numerous slow-type MyBP-C isoforms that result from differential exon splicing (21, 22). Moreover, slow-type MyBP-C is also found in fast-twitch muscles as well, suggesting differential gene expression (23, 24). Therefore, these findings raise the intriguing possibility that a mixture of MyBP-C isoforms may exist in skeletal muscles regardless of twitch speed and their N termini may possess distinct functional capacities to modulate skeletal muscle contractility.

Here, using multi-scale proteomic, biophysical and *in silico* approaches, we define the expression, localization, and modulatory capacities of multiple skeletal MyBP-C isoforms in native skeletal muscle systems. To limit the inherent biological complexity associated with muscle-specific MyBP-C isoform expression (see above), we chose rat slow-twitch soleus (SOL) and fast-twitch extensor digitorum longus (EDL) muscles as model systems. These were the purest rat skeletal muscles with regards to fiber type composition identified in the literature (25). Regardless, we first identified and quantified the abundances and localization of the skeletal myosin and MyBP-C isoforms in the SOL and EDL muscle samples by mass spectrometry and immunofluorescence microscopy. Four different N-terminal MyBP-C isoforms were identified; one being the fast-type MyBP-C isoform with the remaining three slow-type MyBP-C isoforms resulting from alternative N-terminal exon splicing (Fig. 1B). We then determined the Ca^2+^-dependent mechanical interactions between native thin and thick filaments isolated from the SOL and EDL muscles. Thin filament motion was sensitized to Ca^2+^ and maximal sliding velocities were slowed only within the thick filament C-zones. Next, we determined how each of the MyBP-C isoforms contributed to this modulation of actomyosin contractility at the molecular level. We combined *in vitro* motility assays using expressed recombinant N-terminal fragments and *in silico* mechanistic modeling. Our results suggest that each skeletal MyBP-C isoform is functionally distinct, and interestingly, their modulatory capacity depends on the muscle in which they are expressed. Therefore, these rat skeletal muscles use a combination of gene expression and alternative exon splicing in order to fine tune contractility and thus meet these muscles’ diverse physiological demands.

## RESULTS

### Identification and quantification of myosin and MyBP-C isoforms in rat soleus (SOL) and extensor digitorum longus (EDL) muscles by mass spectrometry

To determine muscle myosin and MyBP-C protein compositions, myofibrillar protein preparations were isolated from ~30 mg samples of rat slow-twitch SOL and fast-twitch EDL skeletal muscles (see Methods). We digested these protein preparations with trypsin, analyzed the peptides by liquid chromatography mass spectrometry (LCMS) and quantified protein abundances by label-free analyses (26, 27). Unique peptides generated from the digestion of five myosin heavy chain isoforms were present at varying levels in the myofibrillar protein samples (Table 1). Slow-myosin heavy chain expressed from the *MYH7* gene in the SOL samples and fast-myosin heavy chains expressed from the *MYH4* and *MYH1* genes in the EDL samples predominated (Table 1). The presence of multiple myosin heavy chain isoforms in both the rat SOL and EDL samples confirmed these muscles were composed of mixed fiber-types at the proportions previously reported (28, 29).

**Table 1.**
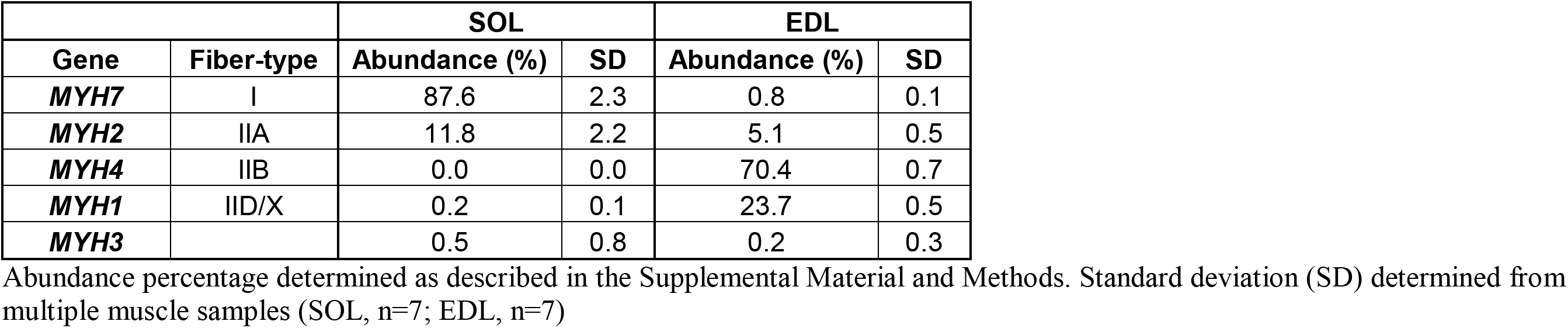
Myosin heavy chain isoform abundance in rat slow-twitch soleus (SOL) and fast-twitch extensor digitorum longus (EDL) skeletal muscles determined by mass spectrometry.

|  |  | SOL |  | EDL |  |
| --- | --- | --- | --- | --- | --- |
| Gene | Fiber-type | Abundance (%) | SD | Abundance (%) | SD |
| *MYH7* | I | 87.6 | 2.3 | 0.8 | 0.1 |
| *MYH2* | IIA | 11.8 | 2.2 | 5.1 | 0.5 |
| *MYH4* | IIB | 0.0 | 0.0 | 70.4 | 0.7 |
| *MYH1* | IID/X | 0.2 | 0.1 | 23.7 | 0.5 |
| *MYH3* |  | 0.5 | 0.8 | 0.2 | 0.3 |

Abundance percentage determined as described in the Supplemental Material and Methods. Standard deviation (SD) determined from multiple muscle samples (SOL, n=7; EDL, n=7)

To determine both the overall content and specific MyBP-C isoforms expressed in these muscles, we analyzed peptides originating from the tryptic digestion of the various MyBP-C isoforms (Fig. S1). The total amount of MyBP-C relative to myosin heavy chain for each muscle was determined based on the abundance of a single, common peptide shared among all MyBP-C isoforms identified (Fig. S1), relative to that of peptides shared among the myosin heavy chain isoforms expressed in that muscle. In the SOL muscle, we estimate 1 MyBP-C molecule per 11.4 ± 0.7 (SEM, n=7) myosin heavy chain molecules, which is no different than the EDL with a ratio of 1 to 11.2 ± 0.6 (SEM, n=7).

Unique peptides associated with multiple slow-type MyBP-C isoforms (*MYBPC1* gene) were present in both the SOL and EDL samples (Fig. S1). In contrast, unique peptides associated with fast-type MyBP-C (*MYBPC2* gene) were present only in the EDL samples (Fig. S1). These relative abundances of the slow and fast-type peptides were indicative of 43.1 ± 3.9% (SEM, n=7) expression of fast-type MyBP-C in the EDL samples. Next, in a more focused analysis of the MyBP-C isoform composition, we enhanced the detection of the various MyBP-C peptides by separating MyBP-C from the other myofibrillar proteins on SDS polyacrylamide gels and then trypsin-digested the 75-150 kDa gel region in preparation for mass spectrometry (n=6 samples per group). Several unique peptides identified in these analyses were indicative of alternative *MYBPC1* gene splicing resulting in both C- and N-terminal slow-type MyBP-C variants. The C-terminal splice variants were found only in the SOL samples (Fig. S1). With the total MyBP-C content relative to myosin heavy chain content in the SOL and EDL samples being similar (see above), the slow-type MyBP-C C-terminal variants do not appear to affect the total number of MyBP-C molecules incorporated into the backbone of the thick filament.

Several peptides in both the SOL and EDL samples were indicative of alternative gene splicing in the 5’ region of the *MYBPC1* gene, resulting in N-terminal slow-type MyBP-C variants (Fig. 1B). The longest of these variants (sMyBP-C) contained amino acids from all N-terminal exons (Table 2, Figs. 1B, S1, S2). A shorter variant lacked exons 4 and 5 (amino acids 35-59, denoted sMyBP-C_Δexon4-5_) (Table 2, Figs. 1B, S1, S2). Interestingly, the combined abundances of these N-terminal peptides associated with the sMyBP-C and sMyBP-C_Δexon4-5_ variants was consistently lower than the abundances of C-terminal peptides. This finding was consistent with the 40% expression of a third short N-terminal variant that uses an alternate start codon in the middle of exon 7, as described by Ackermann et al. (22). This variant lacks the first 125 amino acids originating from the first 6.5 exons (sMyBP-C_Δexon1-6.5_, Table 2, Figs. 1B, S2), beginning midway through the C1 Ig-like domain (Figs. 1B, S2) and based on its sequence would not be identified in our MS analyses. Therefore, the abundance of this variant was assumed to be that reported by Ackermann et al. (22) (Table 2), and then used to estimate by a mass-balance approach (26, 27) the relative abundances of the two longer N-terminal variants (Table 2). Collectively, these data demonstrate that both slow- and fast-twitch muscles utilize alternative exon splicing of the *MYBPC1* gene to modify the structure and possibly the function of slow-type MyBP-C.

**Table 2.**
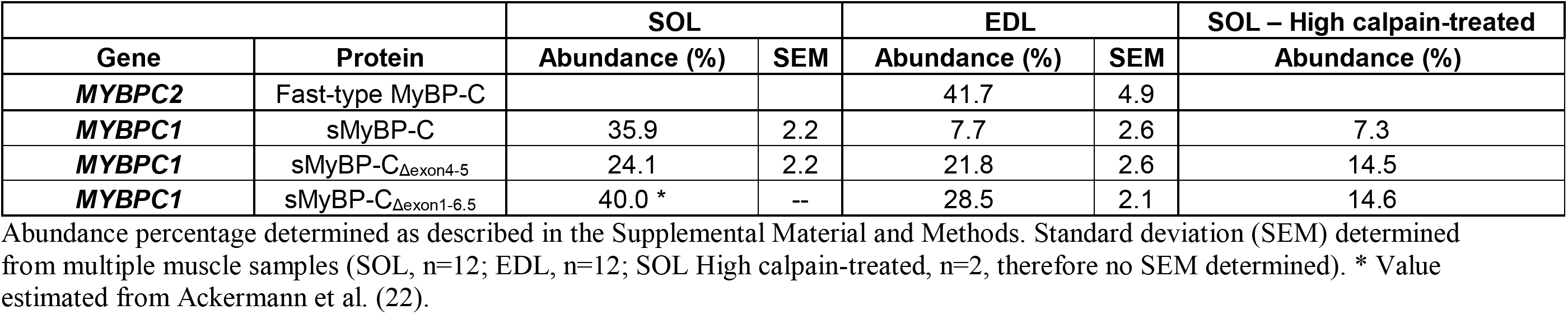
Slow-type and fast-type MyBP-C isoform abundance in rat slow-twitch soleus (SOL) and fast-twitch extensor digitorum longus (EDL) skeletal muscles.

### Spatial distribution of slow- and fast-type MyBP-C in the sarcomere

To determine if MyBP-C isoforms are differentially localized within the C-zones, we immunofluorescently labeled MyBP-C in cryo-sectioned SOL and EDL muscle samples, and imaged them using confocal microscopy (Fig. 2, inserts). The slow-type antibody, capable of binding all slow-type MyBP-C isoforms, labeled both the EDL and SOL muscle samples, while the fast-type MyBP-C antibody only labeled the EDL muscle samples. This was expected from the mass spectrometry data. Both antibodies resulted in two closely spaced, fluorescence bands in the sarcomere center; identifying the two C-zones (Fig. 2). To determine the spatial distribution of the slow- and fast-type MyBP-C isoforms within the C-zones, the signal to noise ratio of the fluorescence signal was improved by averaging multiple sarcomeres (see Methods). The two fluorescent bands were fitted by the sum of two Gaussians, with the fits serving to constrain an analytical model that predicted the spatial distribution of slow- and fast-type MyBP-C molecules within each C-zone (See Methods and Supplemental Information for details).

**Figure 2.**
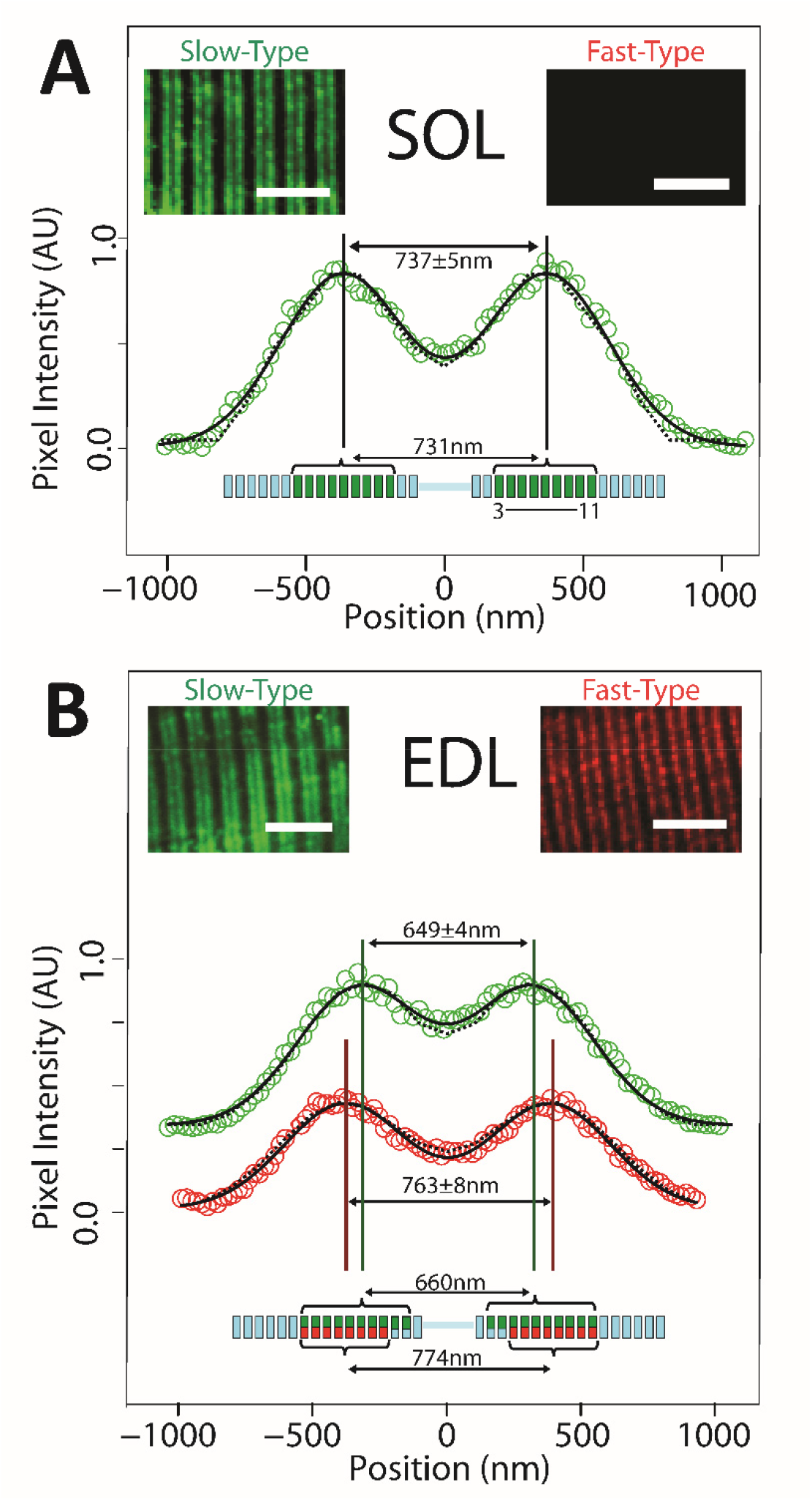
Immunofluorescence imaging and modeling of MyBP-C distribution in SOL and EDL muscle sections. **(A)** In slow-twitch SOL muscle, anti-slow-type MyBP-C antibodies label two bands in each sarcomere. The aligned and integrated intensity of these two bands (open circles, see Supplemental Methods) are well fit with to Gaussian peaks (solid line) with a separation of 737 ± 4.6 nm. Analytical modeling (dotted line, see Supplemental Methods) demonstrated that if slow-type MyBP-C is distributed across thick filament stripes 3-11 (green stripes in thick filament cartoon below), the anticipated peak-to-peak separation would be 731 nm, coincident with the immunofluorescence data. Fast-type MyBP-C was not detected in SOL muscle. **(B)** For fast-twitch EDL muscle, doublet bands were apparent upon labeling with either slow-type or fast-type MyBP-C antibodies. Intriguingly, the separation distance of the two fitted Gaussians peaks differed for slow-type and fast-type MyBP-C. The spatial distribution for slow-type MyBP-C (649 ± 3.9 nm peak-to-peak separation, green) is closer towards the sarcomere midline than fast-type MyBP-C (763 ± 7.6 nm separation, red). Our model suggests that slow-type and fast-type MyBP-C co-occupy stripes 4-11 (green and red stripes, respectively, in thick filament cartoon below data), slow-type MyBP-C may also occupy stripes 2 and 3, although at reduced occupancy (i.e. 2 of 3 possible MyBP-C molecules per stripe, indicated with partially green stripes in thick filament cartoon). This occupancy pattern would result in 660nm peak-to-peak spacing for slow-type MyBP-C and 774nm for fast-type MyBP-C.

To determine how the MyBP-C molecules are distributed within the C-zones of the SOL sarcomeres (Fig. 2A), we developed an analytical model that generated a dual Gaussian fluorescence image based on the point-spread function of the antibody’s fluorophores and their localization along the thick filament (see Supplemental Information for model details). The model assumes that MyBP-C molecules can be located in any of 17 stripes along each half of the thick filament (5, 14). These stripes are associated with the myosin helical repeat and are separated by 43 nm (Fig. 2, schematic). Based on there being 300 myosin heavy chain molecules per half thick filament and our LCMS ratio of 1:11.4 slow-type MyBP-C molecules per myosin heavy chain in the SOL muscle samples, the model assumes there are 27 slow-type MyBP-C molecules per half thick filament. The model then distributes these 27 MyBP-C molecules among the 17 stripes until a MyBP-C distribution is achieved for which the model generated, dual Gaussian fluorescent image best matches the experimental data (Fig. 2A) (Kolmogorov–Smirnov test, p = 0.86, where p > 0.01 demonstrates significant overlap). The best fit was generated by 3 MyBP-C molecules in each of 9 stripes between stripes 3-11 (Fig. 2A). This localization was nearly identical to that measured for rabbit SOL muscle by immuno-electron microscopy (5, 14).

Similar image analysis and modeling was performed on the EDL muscle samples. With both slow- and fast-type MyBP-C isoforms in the EDL, the model assumed that the 27 MyBP-C molecules per half thick filament were divided into 16 slow- and 11 fast-type, based on the LCMS data (Table 2). The best fit distribution of these MyBP-C molecules (p = 0.83) suggests that the slow-type MyBP-C molecules in the EDL samples occupy stripes 2-11 (Fig. 2B, green). Whereas, the fast-type MyBP-C molecules were best distributed (p = 0.94) between stripes 4-11 (Fig. 2B, red). Where the slow- and fast-type MyBP-C molecules occupy the same stripe, a total of 3 MyBP-C molecules are assumed to be present. Therefore, the C-zone spatial distribution of MyBP-C appears to be both MyBP-C isoform and muscle-type specific.

### MyBP-C isoforms differentially modulate native thin filament motility in native thick filament C-zones

We first set out to determine whether the endogenous complement of MyBP-C isoforms in the C-zones of thick filaments can modulate thin filament motility in a Ca^2+^-dependent manner. Therefore, we isolated native thin and thick filaments from both rat SOL and EDL muscles and observed the motility of Ca^2+^-regulated thin filaments over their corresponding native thick filaments to maintain physiological relevance.

#### Thin filament motility is both sensitized to Ca^2+^ and slowed in thick filament C-zones

Native thick filaments present a unique opportunity to characterize how MyBP-C impacts thin filament motility in the C-zone, since the thick filament serves as its own control with its tip being devoid of MyBP-C (i.e. D-zone) (Fig. 3A). Given that the D- and C-zones are each ~350 nm in length (14), we ultrasonically shredded thin filaments into short ~250 nm shards that were capable of interacting within each zone independently. These short thin filaments maintained their Ca^2+^-regulation and sensitivity (Fig. S4) and traversed and sampled both the D- and C-zones (i.e., in the absence and presence of MyBP-C, respectively).

**Figure 3.**
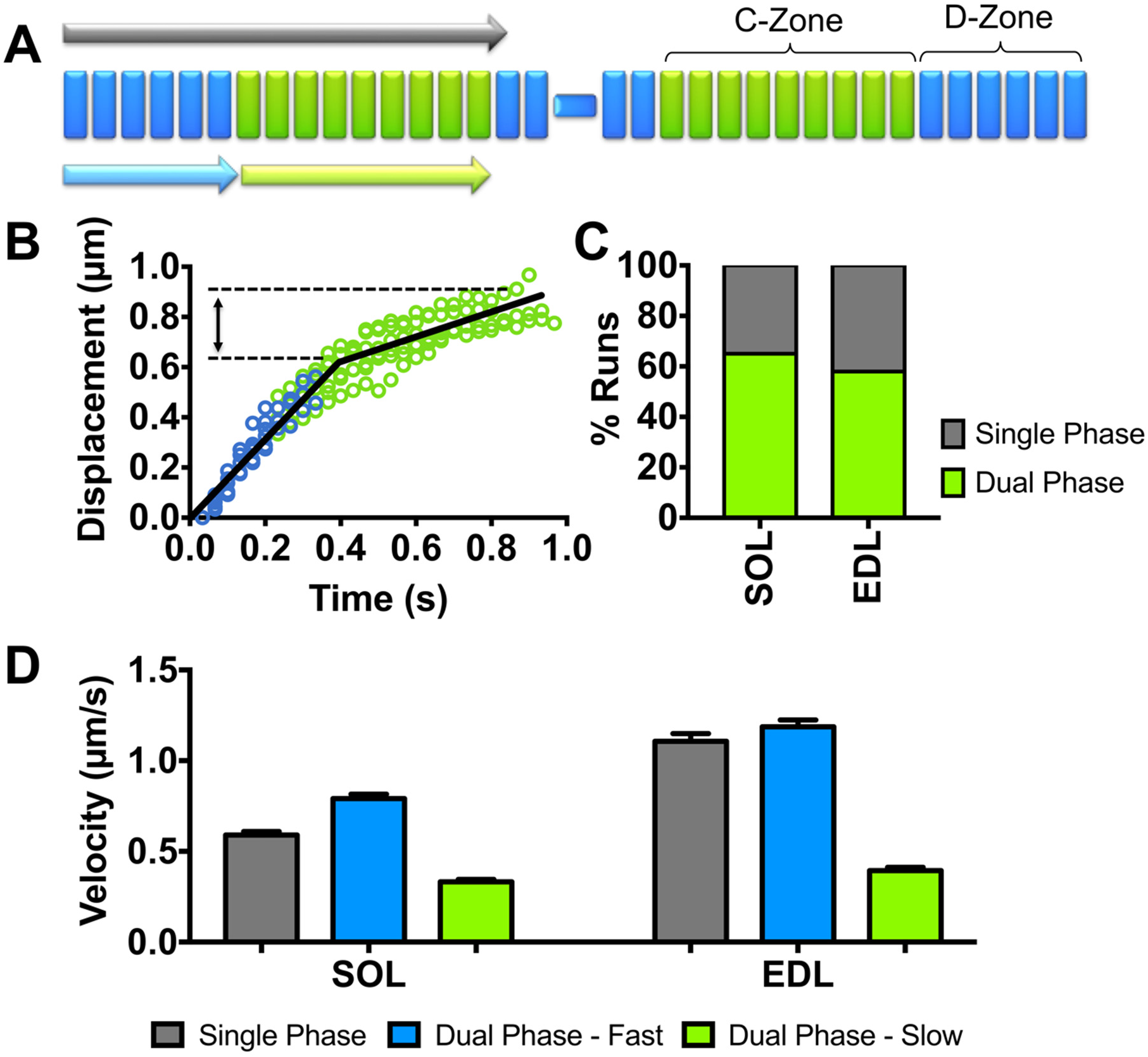
Native thin filament motility at pCa 5 over native thick filaments from SOL and EDL muscles. **(A)** Illustration of native thin filament trajectories observed over native thick filaments, which as in B are described as either a run with a constant velocity (i.e. single phase; grey arrow) or having dual phases, where there is an initial fast velocity phase (light blue arrow) in the D-Zone that is followed by a slower velocity phase (light green arrow) that corresponds to the length of the C-zone. **(B)** Representative EDL thin filament displacement vs. time trajectories illustrating the behavior of dual phase runs. The initial fast velocity phase (blue) persists for ~350-400 nm, approximately the length of the D-zone, followed by a slow velocity phase that corresponds to the expected length of the C-zone (~400 nm, green). **(C)** Percentage of observed thin filament runs that have single versus dual velocity phases. **(D)** The velocities that are associated with single phase and each of the dual velocity phase trajectories for the SOL and EDL thick filaments. The single phase velocities of both SOL and EDL thin filaments are comparable to the fast velocity phase for the dual phase trajectories. All data are presented as mean ± SEM. Data are summarized in Table 3.

At a Ca^2+^ concentration where thin filaments were fully “on” (pCa 5, Fig. S4) as determined in isolated myofibrils (Fig. S4G), the majority (~60%) of trajectories over both SOL and EDL thick filaments showed two distinct phases of velocity (Figs. 3B, 3C). An initial fast velocity phase, with a value correlated to the rate of tension redevelopment in myofibrils and presumably the thick filament myosin type, which then transitioned to a ~60% slower phase, lasting ~400 nm (Figs. 3B-D, Table 3). This distance corresponded to the established C-zone length (30, 31) and the values estimated from our immunofluorescence and modeling data (Fig. 2). The remaining 40% of trajectories over both SOL and EDL thick filaments were better described by a single velocity that was close to the fast velocity phase of the dual-phase trajectories (Figs. 3C, 3D, Table 3). The presence of single-phase velocity trajectories may be due to the inability to identify the velocity transition with statistical certainty or physiological in nature (see modeling below).

**Table 3.** Native thin filament motility over native thick filaments motility under various experimental conditions.

|  |  | pCa 7.5 |  |  | pCa 5 |  |  |  |  |  |  |  | Dual Phase Runs (%) |
| --- | --- | --- | --- | --- | --- | --- | --- | --- | --- | --- | --- | --- | --- |
|  |  |  |  |  | Single Phase Runs |  |  | Dual Phase Runs |  |  |  |  |  |
|  |  | Velocity (μm/sec) | Displacement (μm) | n | Velocity (μm/sec) | Displacement (μm) | n | Fast Phase Velocity (μm/sec) | Fast Phase Displacement (μm) | Slow Phase Velocity (μm/sec) | Slow Phase Displacement (μm) | n |  |
| Low Calpain | SOL | 0.39±0.16 | 0.35±0.14 | 205 | 0.59±0.20 | 0.79±0.04 | 106 | 0.79 ± 0.32 | 0.41±0.15 | 0.33±0.18 | 0.40±0.14 | 197 | 65% |
|  | EDL | 0.46±0.19 | 0.36±0.12 | 135 | 1.11±0.50 | 0.75±0.09 | 133 | 1.19 ± 0.51 | 0.35±0.16 | 0.39±0.25 | 0.47±0.15 | 187 | 58% |
| High Calpain | SOL | NA | NA | NA | 0.57±0.24 | 0.78±0.06 | 172 | 0.72 ± 0.26 | 0.35±0.14 | 0.30±0.13 | 0.50±0.13 | 134 | 44% |

Native thin and thick filaments were isolated using low calpain treatment from rat slow-twitch soleus (SOL) and fast-twitch extensor digitorum longus (EDL) skeletal muscles. The motility of thin filament shards (see Material and Methods) over their respective thick filaments were observed at low $\text{Ca}^{2+}$ (pCa 7.5) and high $\text{Ca}^{2+}$ (pCa 5) concentrations. At pCa 7.5 only short displacements at a constant velocity were observed. At pCa 5, either runs of motility were described by a single velocity and displacement (Single Phase Runs) or by dual phase runs having an initial fast velocity phase followed by a slow velocity phase with the displacements for each phase reported. Following high calpain treatment that proteolytically cleaves only the slow-type MyBP-C N terminus, there was reduction in the percentage of dual phase runs over SOL native thick filaments. n is the number of thin filament runs.

At a Ca^2+^ concentration where thin filaments should have been fully “off” as in myofibrils and thus motility should have been absent (pCa 7.5, Fig. S4), we observed motile thin filaments on both SOL and EDL thick filaments that traveled short, ~350 nm distances at a constant velocity (Fig. 4). Once again, the distances traveled were comparable to the C-zone lengths (Fig. 4B, Table 3), and the velocities were similar to that of the slow velocity phase of trajectories over the SOL and EDL thick filaments at pCa5 (Fig. 3C, Table 3). Therefore, the complement of MyBP-C within the C-zones of both SOL and EDL thick filaments can both sensitize the thin filament to Ca^2+^ at low Ca^2+^ concentrations and slow motility at high Ca^2+^ concentrations.

**Figure 4.**
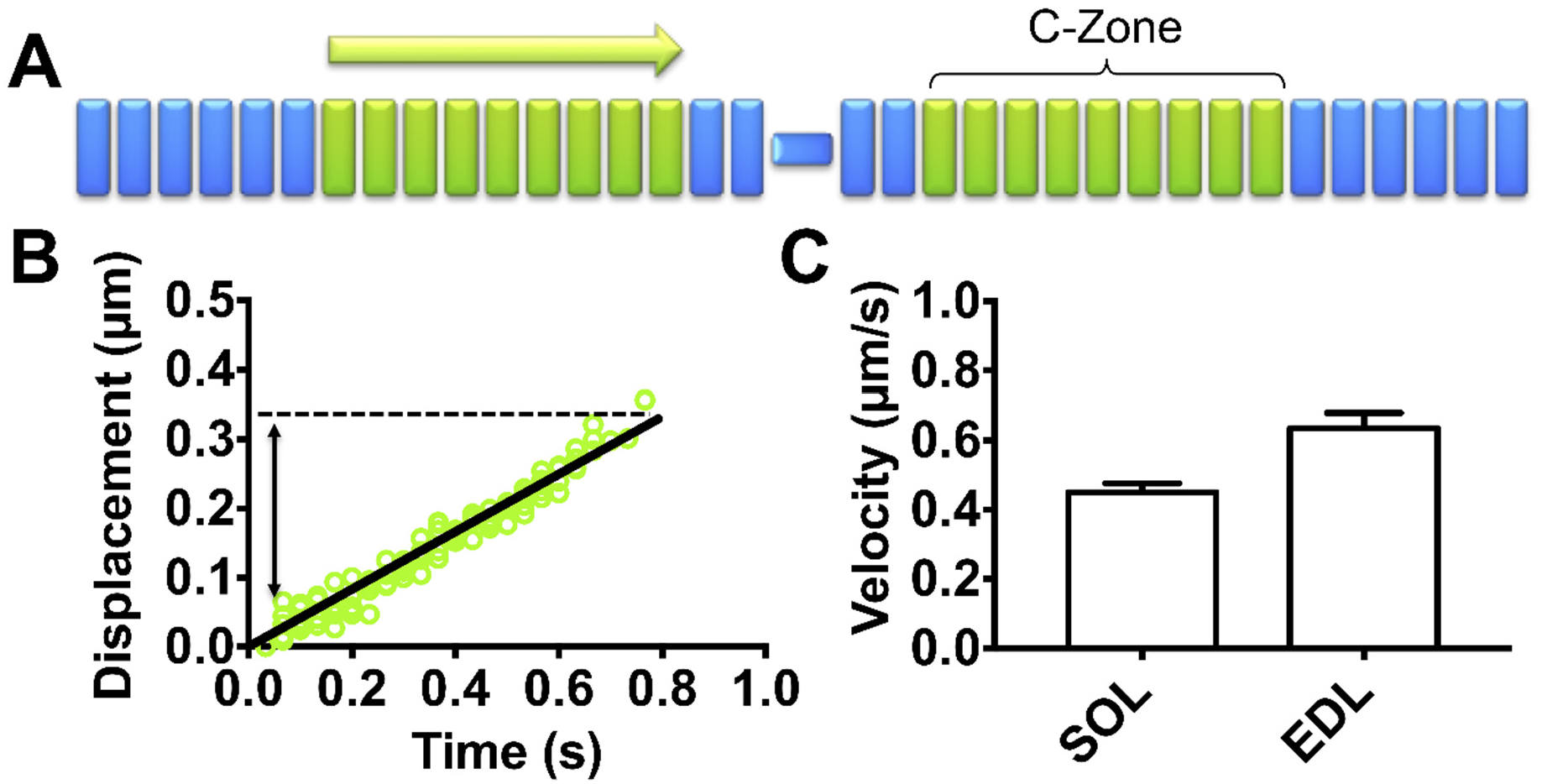
Native thin filament motility at pCa 7.5 over native thick filaments from SOL and EDL muscles. **(A)** Illustration of native thin filament trajectories observed over native thick filaments, which as in B are described by a constant velocity (light green arrow) that corresponds to the length of the C-zone. **(B)** Representative SOL thin filament displacement vs. time trajectories that persist for ~350 nm. **(C)** The velocities of the SOL and EDL thin filament trajectories are comparable to the slow velocity phase of the dual phase trajectories at pCa 5 (Fig. 3D). All data are presented as mean ± SEM. Data are summarized in Table 3.

#### N-terminal domains of slow-type MyBP-C are required to modulate thin filament sliding

To determine if MyBP-C’s N-terminal domains are the effectors of thin filament slowing within the C-zones of native thick filaments, we took advantage of a previously defined thrombin cleavage site in the M-domain that was identified in studies of bovine skeletal MyBP-C (32). Interestingly, this cleavage site target sequence, SAFK, is identical to that for calpain cleavage (27) and exists only in the slow-type MyBP-C variant isoforms (Fig. S2). Therefore, with the SOL thick filaments composed only of the slow-type MyBP-C, we treated these thick filaments with higher calpain concentrations and quantified the effect by LCMS. The abundance of all three slow-type MyBP-C N-terminal variants demonstrated significant reductions in abundance, with the total abundance of slow-type MyBP-C with intact N termini reduced by 64 ± 10% (Table 2).

Following N-terminal cleavage of SOL thick filaments, the proportion of thin filament trajectories at pCa 5 that were characterized by two phases of velocity were no longer the majority, falling from 65% to 44% (Table 3). All of the parameters that described these two-phase velocity trajectories (i.e. fast- and slow-phase velocity values and displacements) were similar to those observed over thick filaments not subjected to high calpain treatment (Fig. 3, Table 3). These results suggest that the N terminus of one or more of the slow-type MyBP-C variant isoforms is responsible for the thin filament slowing observed in the SOL C-zone.

#### Isoform specific N-terminal MyBP-C fragments differentially modulate thin filament motility in a Ca^2+^-dependent manner

To determine the functional contribution of individual MyBP-C N-terminal isoforms, we turned to the conventional *in vitro* motility assay to assess the impact of recombinant N-terminal MyBP-C fragments on SOL and EDL thin filament motility propelled by their respective monomeric myosins. Recombinant N-terminal fragments, up to and including the C2 domain, were bacterially expressed for the fast-type (fC1C2) and the three slow-type N-terminal variants (sC1C2, sC1C2_Δexon4-5_, sC1C2_Δexon1-6.5_) (Fig. 1B). Unfortunately, both bacterial- and baculovirus-expressed sC1C2_Δexon1-6.5_ were insoluble under non-denaturing conditions, thereby precluding their characterization (Fig. S5).

N-terminal fragments were added to the motility assay at pCa 7.5 and pCa 5 to assess each fragment’s ability to sensitize thin filaments to Ca^2+^ and to slow thin filament motility, respectively (Figs. 5, S6). By varying the fragment concentrations (0-2 µM), Hill activation (pCa 7.5) and non-competitive inhibition (pCa 5) curves were generated and fitted to normalized motility data defined by the product of thin filament sliding velocity and the fraction of moving filaments (Figs. 5, S6, Table S1). The product of velocity and fraction moving provides a means of taking into account the activation state of the thin filament (Fig. S4).

**Figure 5.**
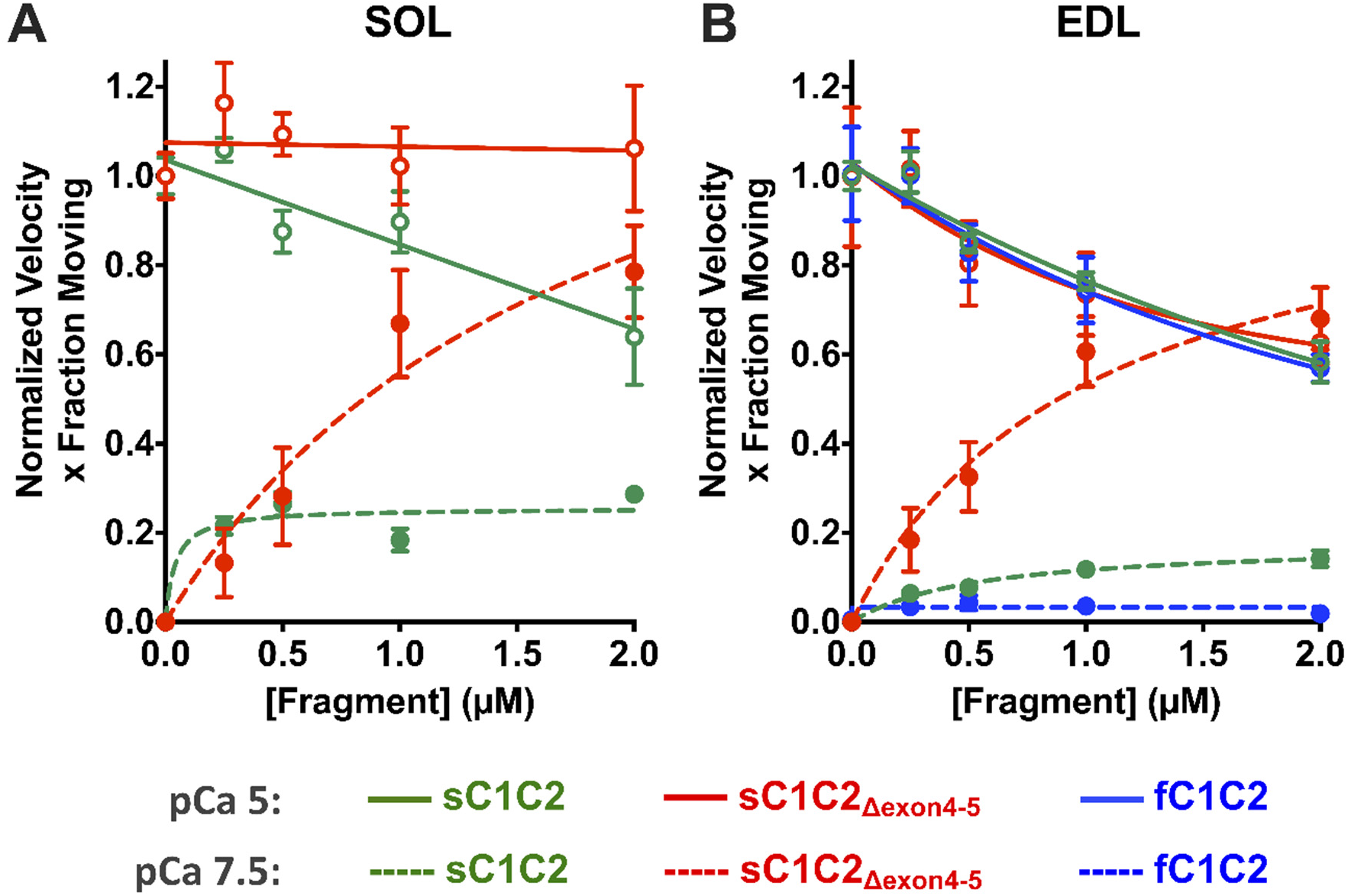
Isoform specific N-terminal MyBP-C fragments differentially modulate SOL and EDL thin filament motility over monomeric myosin in a Ca^2+^-dependent manner. MyBP-C imparted concentration-dependent (0-2µM) effects on thin filament sliding motility at low (pCa 7.5, closed symbols and dashed lines) and high (pCa 5, open symbols and solid lines) calcium concentrations. Plots are shown as velocity times the fraction of filaments moving, normalized to the value at pCa 5 for each MyBP-C fragment. Data were fitted with non-competitive inhibition (K_i_) and Hill activation (K_a_) curves for pCa5 and pCa 7.5, respectively (see Table S1 for K_i_ and K_a_ parameter fits). **(A)** For SOL thin filament motility at pCa 5, sC1C2 reduced motility by up to 36% with increasing fragment concentration (green solid line). sC1C2_Δexon4-5_ had negligible impact on thin filament motility (red solid line). At pCa 7.5, sC1C2_Δexon4-5_ (red dashed line) significantly enhanced thin filament motility with increasing concentrations compared to sC1C2 (green dashed line). **(B)** For EDL motility, all three N-terminal fragments have comparable inhibitory effects on thin filament velocities at pCa 5 (solid lines). At pCa7.5, fC1C2, sC1C2 and sC1C2_Δexon4-5_ differentially sensitize the thin filaments to calcium with sC1C2_Δexon4-5_ (red dashed line) being the greatest sensitizer, compared to sC1C2 (green dashed line) and fC1C2 (blue dashed line). All data are presented as mean ± SEM. See Fig. S6 for plots of individual velocity and fraction of moving filaments and Table S1 for parameters of the fits to the data and the number of experiments for each condition.

For the SOL-based motility at pCa 5, where thin filaments are fully “on”, increasing concentrations of sC1C2 slowed normalized thin filament motility up to 36% with a fragment concentration (K_i_) inhibiting these parameters by 50% of 3.8±1.3 µM (Fig. 5A, green solid line). The shorter, sC1C2_Δexon4-5_ fragment had no effect on motility as evidenced by a K_i_ (120±644 µM), well beyond the 2 µM fragment concentration tested (Fig. 5A, red solid line). For the SOL-based motility at pCa 7.5, where thin filaments were normally “off” (Fig. S4), the shorter sC1C2_Δexon4-5_ fragment had the greatest effect on sensitizing thin filaments to Ca^2+^ (Fig. 5A, red dashed line). At 2 µM sC1C2_Δexon4-5_, thin filament motility was nearly equal to that at pCa 5. In contrast, the sC1C2 was not as effective at sensitizing the thin filaments to Ca^2+^ (Fig. 5A, green dashed line).

In the EDL where both fast- and slow-type MyBP-C isoforms are expressed, we determined the effect of the fC1C2, sC1C2, and sC1C2_Δexon4-5_ N-terminal fragments on thin filament motility in the EDL-based assay at pCa 5. All of the fragments slowed thin filament motility up to 40% with similar K_i_ ~ 2.7 µM (Fig. 5B, solid lines; Table S1). In contrast at pCa7.5, each fragment differentially sensitized thin filament motility to Ca^2+^. Specifically, the sC1C2_Δexon4-5_ fragment was a potent Ca^2+^ sensitizer (Fig. 5B, red dashed line), which was not the case for both the sC1C2 (Fig. 5B, green dashed line) and fC1C2 fragments (Fig. 5B, blue dashed line). Therefore, all of the expressed N-terminal fragments tested, imparted a concentration-dependent effect on sensitizing thin filaments to Ca^2+^ at pCa 7.5 and on slowing motility at pCa 5, however, the extent of these effects were muscle type-specific.

### *In silico* model of thin filament slowing in thick filament C-zones

Knowing how the individual N-terminal fragments modulate thin filament motility as described above, we implemented an *in silico* Monte Carlo-style simulation in an effort to understand how each N-terminal splice variant isoform contributes to the overall slowing of thin filament motility in the C-zones of native thick filaments at pCa 5. Another goal of this modeling effort was to infer the slowing capacity of the sMyBP-C_Δexon1-6.5_ splice variant, which could not be characterized in the *in vitro* motility assay due to the fragment’s insolubility. To simplify these modeling efforts, we focused only on SOL thick filaments due: 1) fewer number of N-terminal isoforms (i.e. 3 for SOL vs. 4 for EDL) (Table 2); 2) profound difference in the slowing capacity for the characterized fragments (Fig. 5A), and: 3) N-terminal proteolytic cleavage of the slow-type MyBP-C by calpain provided an experimental condition in which a proportion of the existing MyBP-C were no longer functional.

The model and its simplifying assumptions are described in detail within the Supplemental Information but briefly are as follows:

1. Each SOL half thick filament is comprised of seventeen “stripes” spaced 43 nm apart as defined by the myosin helical repeat (Fig. 6A). Each stripe generates a maximal velocity of 0.79±0.32 μm/s that equals the experimentally observed fast velocity phase of a thin filament trajectory (Fig. 5, Table 3);
2. The slow-type MyBP-C N-terminal splice variant isoforms are randomly distributed in stripes 3-11 of the C-zone as determined by our immuno-histological imaging (Fig. 2), with the probability of a splice variant occupying a stripe determined by the LCMS abundance data (Table 2).
3. Each C-zone stripe has an inherent velocity that is slower than the maximal velocity based on the inhibition constant (K_i_) (Table S1, Fig. 5) for the splice variant assigned to that stripe.
4. A short thin filament (250±50 nm) initiates a trajectory near the thick filament tip (stripe 17) and proceeds through the C-zone (Fig. 6A). As the thin filament is translocated over the thick filament, its effective velocity at any point is the mean of the velocities associated with the ~6 stripes that the thin filament is in contact with. Therefore, the thin filament velocity is position-dependent.
5. To simulate the experimental condition following high calpain treatment, N-terminally cleaved slow-type MyBP-C are assumed to no longer slow velocity. Therefore, the probability of these cleaved splice variants being in a C-zone stripe is determined by their relative abundance, measured by LCMS (Table 2). The stripe in which they are assigned thus, generates the uninhibited maximal velocity.
6. Simulated trajectories (i.e. 1,000) were parameterized to match the experimental frame rate (30 fps) and spatial resolution (14 nm) and analyzed for the number of velocity phases by the same approach employed experimentally (see Supplemental Information for details). Once analyzed, velocity distributions for trajectories with a single constant velocity and for the fast and slow phases of the dual-phase trajectories were generated and the model distributions then compared to the experimentally obtained distributions (Figs. 6D, 6E).
7. The model has only two free parameters: 1) the effective total slow-type MyBP-C concentration; 2) the inhibition constant (K_i_) for the uncharacterized slow-type MyBP-C_Δexon1-6.5_ splice variant. Therefore, trajectories were simulated for two experimental conditions (i.e. low and high N-terminal calpain cleavage) over a range of possible values for the two free parameters (Fig. 6B). For each condition and free parameter value, the simulated velocity distributions and fraction of trajectories that were one versus two phases were compared to the experimental data and assigned a “goodness of fit” (see Supplemental Information). The goodness of fit, over the entire parameter space explored, is presented as a color heat map (Fig. 6B). Using the best fit parameters (4 µM total [MyBP-C], 26 µM K_i_ for MyBP-C_Δexon1-6.5_), the model effectively recapitulates the trajectory characteristics before and after N-terminal cleavage with respect to the proportion of dual-phase trajectories (Figs. 6D, 6E inset) and the velocity distributions for the fast- and slow-phase of the trajectory (Figs. 6D, 6E). The model suggests that the ~40% of trajectories that have a single velocity phase can in part be due to the stochastic assignment of slow-type MyBP-C isoforms to the various stripes (Fig. 6A). Specifically, if the N-terminal variants that are minimally inhibitory to thin filament sliding velocity populate the stripes closest to the D- to C-zone transition, then the MyBP-C slowing effect would not occur until the thin filament is well into the C-zone. This would compromise our ability to statically detect the transition between the fast and slow velocity phases of the trajectory. Although the simulation did not return a single unique parameter set, it did identify a range of parameters that gave equally good fits. Specifically, the effective total slow-type MyBP-C concentration was predicted to be 3-12 μM. This is in agreement with an ~11 μM estimate, based on our LCMS ratio of slow-type MyBP-C to myosin heavy chain of 1:11.4 (Table 1 and 2), assuming a myosin concentration of ~120 μM in skeletal muscle (33). More importantly, the estimated K_i_ for the uncharacterized slow-type MyBP-C_Δexon1-6.5_ N-terminal variant isoform is no less than 25µM (Fig. 6B), which would suggest that this isoform may have negligible impact on thin filament sliding velocity in the C-zone.

**Figure 6.**
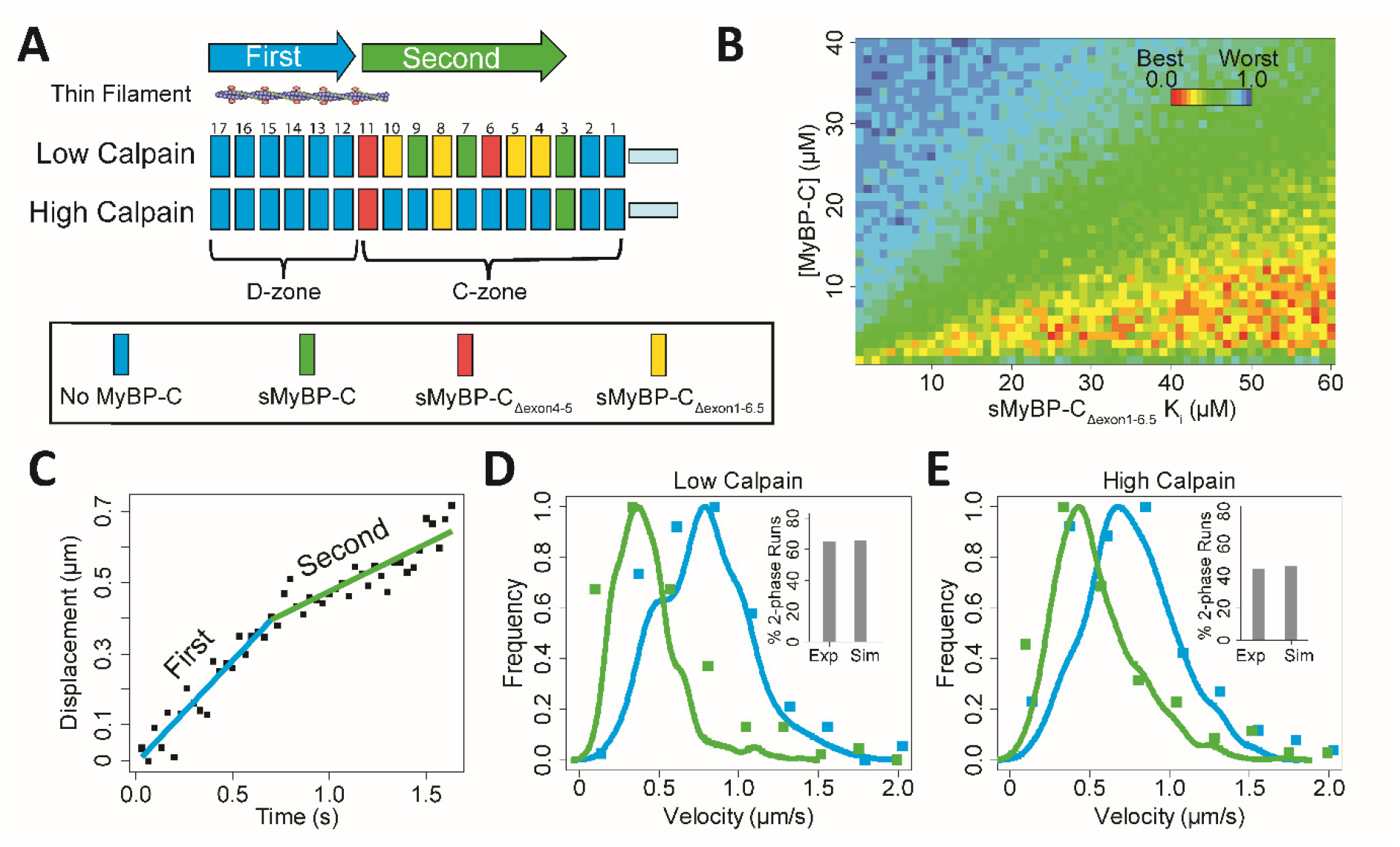
*In silico* modeling showing the slowing effects on thin filament motility at pCa 5 of the slow-type MyBP-C isoform complement in the C-zones of native SOL thick filaments. In addition, the model was used to extrapolate the functional influence of sMyBP-C_Δexon1-6.5_, which could not be characterized by *in vitro* motility (see main text and Supplemental Information for model details). **(A)** Schematic illustrating the localization of slow-type MyBP-C molecules in stripes 3-11 within the C-zone of native thick filaments. The probability that a stripe is occupied by a specific MyBP-C isoform is related to the relative abundance for the isoform as determined by LCMS (Table 2). At low calpain concentrations, slow-type MyBP-C N termini remain intact while high calpain treatment cleaves 64% the N termini (Table 2), thereby reducing the intact MyBP-C stripe occupancy down to ~3 stripes within the C-zone. **(B)** The model has two variable parameters, the effective total MyBP-C concentration and the K_i_ (inhibition constant) for the sMyBP-C_Δexon1-6.5_ isoform. The heat-map shows the normalized goodness of fit for a range of modeled parameters that are used to generate model thin filament trajectories as compared to the experimental data from Fig. 3 and Table 3. There is no unique best fit parameter set but rather a range of best fit parameters for the effective total slow-type MyBP-C concentration of 3-12 µM. More importantly, the predicted K_i_ being >25µM for the sMyBP-C_Δexon1-6.5_ isoform, which is well above the effective total MyBP-C concentration, suggests this isoform has little to no inhibitory effect on thin filament slowing. (**C**) Displacement vs. time trajectories generated by the model, showing the dual velocity phases. (**D**) Velocity distributions for the individual phases of the dual velocity phase trajectories (fast velocity phase, blue; slow velocity phase, green) for the simulated data (solid lines) compared to the experimental data (solid symbols), with the percentage of dual phase runs for the experimental (Exp) and simulated (Sim) data in the insets. (**E**) As in D but after high calpain treatment. All plots demonstrate that the simulated data generated using the modeled best-fit values (4 µM total [MyBP-C], 26 µM K_i_ for MyBP-C_Δexon1-6.5_) closely resembles experimental native thick filament data.

## DISCUSSION

Using multi-scale proteomic, biophysical and *in silico* approaches, we defined the expression, localization, and modulatory capacities of MyBP-C isoforms in native myosin thick filaments from rat slow-twitch SOL and fast-twitch EDL skeletal muscle. Specifically, in SOL at least three slow-type MyBP-C isoforms are expressed that differ in their N termini as a result of alternate exon splicing (Figs. 1B, S2, Table 2). Whereas, in the EDL, these same slow-type MyBP-C isoforms are expressed with the addition of a fast-type MyBP-C (Figs. 1B, S2, Table 2). In native thick filaments, this mixed complement of MyBP-C isoforms both sensitizes the thin filament to Ca^2+^ while slowing thin filament velocity, only within the C-zones where MyBP-C is localized (Figs. 3, 4). The major effector of this modulatory effect on filament sliding is the MyBP-C N terminus (Fig. 5, Table 3, Table S1), which presumably projects away from the thick filament backbone and interacts with the thin filament and/or the myosin head region (14).

Although collectively, the mixed population of MyBP-C isoforms in the C-zone modulates thin filament motility, how each individual isoform contributes to these effects is far more complex. For example, in the EDL thick filament C-zone, where both slow- and fast-type MyBP-C isoforms are present, are these proteins spatially segregated? Are the fast-type and slow-type MyBP-C and its variants functionally identical? If not, then are functional differences dependent on the muscle-specific thick filament in which they exist? Only through the use of multi-scale biophysical approaches described here, have we been able to address these questions.

### Expression and spatial distribution of MyBP-C isoforms in the C-zone

The expression of slow-type MyBP-C isoforms in muscle has been observed in both slow twitch, type-I and fast twitch, type II muscle fibers (22). Whereas, fast-type MyBP-C is solely expressed in fast twitch, type II muscle fibers (20). These general observations agree with our immunofluorescence data, where the slow-type MyBP-C is detected both in the rat slow-twitch SOL and fast-twitch EDL muscles, while the fast-type MyBP-C is only detected in the EDL (Fig. 2, Table 2). However, fast twitch, type II muscle fibers can be further categorized by their myosin heavy chain isoform expression (i.e. type IIA, IIB, IIX). Therefore, the presence of as much as 20% fast twitch, type IIA myosin in the rat SOL, based on our own LCMS data (Table 1) and the literature (29), suggests that the assumed expression of fast-type MyBP-C in all fast twitch, type II fibers may not hold. Specifically, the absence of any fast-type MyBP-C in the rat SOL (Table 2), even though fast twitch, type IIA fibers do exist, suggests that the type IIA fibers in the SOL only express slow-type MyBP-C. Whether this applies to the type IIA fibers in the rat EDL, which only make up 5% of the fiber composition (Table 1), cannot be determined based on the existing data. Interestingly, type IIA fibers are the slowest of the fast twitch fibers and thus closer in speed to type I fibers (34). This might imply that a muscle fiber’s speed dictates higher order transcriptional or translational regulation that determines the pattern of MyBP-C isoform expression.

Regardless of the MyBP-C isoform expression pattern, how are the slow- and fast-type MyBP-C isoforms spatially distributed in the C-zones of SOL and EDL muscle? Evidence from immuno-electron micrographs (5, 14) localize MyBP-C within the C-zone to 7-9 transverse stripes separated by 43 nm (Figs. 1A, 2). In agreement with the literature, using a spatially explicit analytical model to help interpret our immunofluorescence data from tissue samples, the slow-type MyBP-C in the SOL appears to distribute into 9 stripes within the C-zone (Fig. 2A). Since the antibody epitope cannot distinguish between the various slow-type isoforms, the specific distribution of these isoforms amongst the 9 stripes cannot be determined. However, in the EDL where both slow- and fast-type MyBP-C are expressed, the model does predict that these two isoforms are spatially distinct. Even without modeling, the raw immunofluorescence data imply that the peaks of the slow-type MyBP-C doublets compared to that of the fast-type MyBP-C are on average ~60 nm closer to the sarcomere center (i.e. M-line) (Fig. 2B). More precisely, the model suggests that the fast-type MyBP-C is distributed into 8 stripes (stripes 4-11) within the C-zone, whereas the slow-type MyBP-C is distributed into 10 stripes (stripes 2-11). This implies that the stripes closest to the M-line in the EDL are only occupied by the slow-type MyBP-C. What governs this spatial segregation in the C-zone, not to mention the stoichiometry of the various MyBP-C isoforms? Regulated expression could be one factor that determines stoichiometry. More interestingly, isoform differences in the MyBP-C C-terminal C10 domain, which occurs by alternate exon splicing in the slow-type MyBP-C (see Results, Figs. S1, S3), were observed here and by others (22). Although these differences do not affect the overall stoichiometry, they may dictate an isoform’s binding affinity to particular locations along the thick filament through interactions with the backbone itself and/or titin (35). Regardless, the observed difference in the overall spatial distribution of MyBP-C isoforms in the C-zone might be related to functional differences between the isoforms, as described below.

### Skeletal MyBP-C isoforms modulate thin filament motility

We demonstrated that the N-terminal domains of MyBP-C are the effector of MyBP-C’s actomyosin modulatory capacity (Fig. 5, Table 3). Therefore, we used bacterially expressed N-terminal fragments (up to and including the C2 Ig-domain) of the various slow- and fast-type MyBP-C isoforms in a motility assay to define each isoform’s potential contribution to thin filament Ca^2+^-sensitization and slowing of velocity (Fig. 5). Based on the endogenous MyBP-C profile (Table 2), we expressed the fast-type MyBP-C N-terminal fragment (fC1C2) and the three slow-type MyBP-C N-terminal splice variants (sC1C2, sC1C2_Δexon4-5_, sC1C2_Δexon1-6.5_) (Fig. 1B).

#### Isoform differences in thin filament Ca^2+^-sensitization

In the absence of MyBP-C, Ca^2+^-regulated thin filaments are effectively “off” at pCa 7.5 (Fig. S4), but on native thick filaments from either SOL or EDL muscle samples, thin filament motility is apparent in the C-zone (Fig. 4). This is presumably due to MyBP-C’s Ca^2+^-sensitizing capacity. When the various N-terminal fragments were characterized in the conventional motility assay, the sC1C2_Δexon4-5_ fragment was the most effective Ca^2+^-sensitizer in both SOL and EDL-based motility assays (Fig. 5A, 5B, red dashed line), whereas the other fragments displayed more modest Ca^2+^-sensitization (Fig. 5, green and blue dashed lines). Therefore, the sC1C2_Δexon4-5_ splice variant isoform may be the major contributor to thin filament Ca^2+^-sensitization in both the SOL and EDL thick filament C-zones (Fig. 4). In a previous study, we investigated how heterologous expression of skeletal MyBP-C isoforms in cardiac muscle during heart failure (18) might impact MyBP-C’s modulation of cardiac contractility. Interestingly, even within the context of a cardiac contractile protein system (i.e. cardiac myosin and thin filaments), the sC1C2_Δexon4-5_ fragment (referred to as ssC1C2 in that study) was more effective at Ca^2+^-sensitizing cardiac thin filaments than the fC1C2, suggesting that the enhanced Ca^2+^-sensitization by the sC1C2_Δexon4-5_ fragment is not muscle specific (18). One mechanism by which the sC1C2_Δexon4-5_ fragment may Ca^2+^-sensitize the thin filament, as was the case for this fragment in the cardiac system (18), is its binding to the thin filament to shift tropomyosin from the “blocked” to the “closed” state, which even at pCa 7.5 would then allow myosin to interact with the thin filament (36–38). From a structural standpoint, both the slow- and fast-type MyBP-C can Ca^2+^-sensitize the thin filament to varying degrees in the motility assay, but do so without a C0 Ig-like domain that is present in cardiac MyBP-C. The lack of this extra C0 domain may limit the extent to which the slow- and fast-type MyBP-Cs sensitize the thin filament to Ca^2+^. Structural studies demonstrate that the C0 domain by itself can bind to the thin filament, but is incapable of activating the thin filament (36). Therefore, the slow- and fast-type MyBP-C lacking the C0 domain would not have this additional weak point of contact with the thin filament, which if present may increase the probability that the other N-terminal domains stereospecifically interact with the thin filament and shift tropomyosin to a position more favorable for myosin binding.

#### Isoform differences in slowing of thin filament velocity

At high Ca^2+^ concentrations (pCa 5), where thin filaments are effectively “on” (Fig. S4), thin filament velocities slowed in both the SOL and EDL thick filament C-zones, thus defining MyBP-C’s second modulatory capacity (Figs. 3, 5). This thin filament slowing upon entering the C-zone has recently been confirmed during fiber shortening in skinned rat soleus skeletal muscle fibers (19). Thin filament slowing appears to be inherent to the N termini of all slow- and fast-type MyBP-C fragments characterized in the motility assay with two exceptions (Fig. 5, solid lines). The sC1C2_Δexon1-6.5_ variant, which could not be characterized due to its insolubility, was predicted to have no mechanical impact based on our *in silico* modeling efforts (see Results). If so, this may not be surprising since this variant’s sequence starts halfway through the C1 Ig-like domain (Fig. S2). Previous studies with mouse cardiac N-terminal fragments showed that any fragment with slowing capacity required an intact C1 domain (11, 36). The other exception is the sC1C2_Δexon4-5_ variant, which is inhibitory in the EDL-based motility assay (Fig. 5B, red solid line) but not in the SOL-based motility assay (Fig. 5A, red solid line). Therefore, the observed slowing is context-specific in the sense that the same sC1C2_Δexon4-5_ N-terminal variant functions differently depending on the thin and thick filament proteins it interacts with, i.e. SOL versus EDL contractile proteins. Although Ackermann et al. (17) reported different degrees of bare actin filament slowing for several recombinant mouse slow-type MyBP-C N-terminal splice variant fragments (up to and including the M-domain), their motility assay used actin and myosin from heterologous species and muscle types. Therefore, interpretation of their results must be viewed in the context of our findings in which a MyBP-C fragment’s functional capacity can be dependent on the contractile proteins.

Potential mechanisms that may slow thin filament velocity are that the MyBP-C N-terminal domains act as internal load and/or as a competitive inhibitor that competes with myosin heads for binding to the thin filament. Evidence in the laser trap assay of cardiac MyBP-C N-terminal fragments binding transiently to actin (11) was interpreted as imparting a viscous-like load against which myosin motors must operate (39). Although, MyBP-C N-terminal binding to the myosin head region could also serve as an internal load, restricting myosin head movement and force production (16). Evidence also exists to support MyBP-C competing with myosin for actin-binding based on solution cosedimentation (40), inhibition of ATPase activity (41), and recent direct visualization of reduced binding of S1-myosin to a single thin filament tightrope in the presence of a cardiac N-terminal fragments (42). In fact, Walcott and coworkers combined the viscous load and competitive inhibition modes of MyBP-C interaction with the thin filament into a model that recapitulated the concentration dependence of N-terminal cardiac MyBP-C fragment-induced slowing of actin filament velocity in the motility assay (43). Why the slowing of thin filament velocity for the sC1C2_Δexon4-5_ variant is so different in the SOL-versus the EDL-based motility systems (Fig. 5) is a matter of speculation. For example, subtle differences may exist in the binding interface between this fragment and the protein isoforms that constitute the thin and thick filaments in these two muscle types.

If the shorter slow-type MyBPC_Δexon1-6.5_ N-terminal variant isoform is incapable of slowing thin filament velocity in the C-zone, as suggested by our modeling efforts, then what physiological purpose could it serve? A recent report suggests that in addition to its modulatory role as defined here, slow-type MyBP-C may play a structural role in thick filament assembly within the sarcomere, as knockdown of slow-type MyBP-C in mouse FDB skeletal muscle resulted in reduced abundance of myosin and myomesin (44). Therefore, the 1:11.4 ratio of MyBP-C to myosin, reported here, must be critical to the maintenance of thick filament stability. As such, significant expression of the slow-type MyBPC_Δexon1-6.5_ isoform in both the SOL and EDL thick filaments may allow its intact C-terminal domains to provide structural stability to the thick filament (4).

## Conclusion

MyBP-C appears to sensitize the thin filament to Ca^2+^ while acting as internal “brake” that slows thin filament velocity only within the C-zone. Interestingly, these two modulatory capacities are not common to the multiple MyBP-C isoforms that populate the skeletal muscle C-zone. Rather, the various isoforms can Ca^2+^-sensitize or slow velocity to different extents depending on the muscle fiber type in which they exist. As such, the differential expressions of slow- and fast-type MyBP-C isoforms may enable the muscle to fine-tune its mechanical performance depending on the muscle’s physiological demands. It is worth noting that at least the slow-type MyBP-C is post-translationally modified by phosphorylation of its N-terminal domains (19, 45–47). Therefore, N-terminal phosphorylation of the slow-type MyBP-C may provide a means of regulating its modulatory capacity both normally under acute physiological stress and abnormally due to genetic mutations in MyBP-C that lead to skeletal myopathies (19, 45–47).

## Materials and Methods

Recombinant protein isolation, production and motility experiments were performed according to previously published protocols (18, 27). Comprehensive experimental details are provided in the supplementary ‘Material and Methods’-section.

### Proteins

Myosin, native thick filaments and native thin filaments (NTFs) were isolated from soleus (SOL) and extensor digitorium longus (EDL) muscles of Sprague Dawley rats. F-actin was purified from chicken pectoralis muscle (48). Mouse MyBP-C N-terminal fragments (Fig. 1B) up to the C2 domain fC1C2 (1–337), sC1C2 (1–366), sC1C2_Δexon4-5_, (1–341) and sC1C2_Δexon1-6.5_ (1–241) were bacterially expressed as previously described (18) unless stated otherwise.

### Thick filament isolations

Thick Filaments were isolated according to the procedures detailed in Previs et al. (18). Freshly isolated rat SOL or EDL were manually shredded in 0.5mL of Relaxing Buffer (50 mM NaCl, 5 mM MgCl_2_, 2 mM EGTA, 1 mM DTT, 7 mM phosphate buffer (pH 7), 10 mM creatine phosphate, 2.5 mM ATP) over the course of 30 minutes, while kept on ice. Tissue was further dissected in skinning solution (Relaxing Buffer containing 0.5% v/v Triton X-100) over 2 rounds of 30 minutes each, followed by 2 rinses (30 minutes each) with Relaxing Buffer. Finally, thick filaments were liberated by digestion of a small amount of skinned fibers over several minutes at room temperature, using Calpain-1 from porcine erythrocytes (Calbiochem), at either 0.05U/uL (Low Calpain) or 0.25U/uL (High Calpain) concentrations in buffer containing 1mM calcium acetate, 20mM imidazole (pH 6.8), and 5μM 2-mercaptoethanol. The slide was rinsed repeatedly with 10-μl aliquots of Relaxing Solution (supplemented to 75 mM NaCl total), and all washes were combined in a microcentrifuge tube. The total volume of the tube was raised to 250μl and the contents were clarified by centrifugation at 1,000g for 1 minute.

### Quantitative mass spectrometry

Briefly, thick filaments samples were isolated and digested in solution with trypsin, or MyBP-C was separated by SDS-PAGE and digested in-gel, as described in the Results. Aliquots of each sample, were separated by ultra-high pressure liquid chromatography and the eluent was analyzed using a Q Exactive Hybrid Quadrupole-Orbitrap mass spectrometer (Thermo Fisher Scientific). Peptides were identified from the resultant spectra using SEQUEST in the Proteome Discoverer 2.2 (PD 2.2) software package (Thermo Fisher Scientific) and custom-built databases. Quantification was carried out using a label-free approach (26, 27). Details are described in the supplemental Material and Methods.

### Native thin filament *in vitro* motility

Conventional *in vitro* motility assays were performed as previously described in (18). Briefly, myosin was incubated onto a nitrocellulose coated surface of a glass flow cell. Rhodamine labeled NTFs were incubated in the flow cell prior to the addition of bacterially expressed fragments and activating buffer. The motion of NTFs were observed by epifluorescence microscopy. Myosin and NTFs were muscle-type matched in all experiments.

### Native thick filament *in vitro* motility

Native thick filament *in vitro* motility assays were performed as detailed previously (27). Sigmacote (Sigma Aldrich)-treated flow-cells were initially incubated with freshly prepared native thick filaments, and allowed to incubate at room temperature for 20 minutes. The flow-cell surface was then blocked with BSA (1mg/mL). This was then followed by rhodamine labeled thin filaments, which were sonicated (Model 300, Fisher) immediately prior to addition to the flowcell. Finally, motility buffer (25 mM KCl, 1 mM EGTA, 10 mM DTT, 25 mM imidazole (pH 7.4), 4 mM MgCl_2_, 100μM ATP) with an oxygen scavenging system (0.1μg/mL glucose oxidase, 0.018μg/mL catalase, 2.3μg/mL glucose) with freshly sonicated thin filaments and at the indicated pCa is introduced into the flowcell. The motion of the NTFs were captured under total internal reflection fluorescence imaging at 30FPS. Importantly, these motility experiments were performed with native thick filaments and NTFs derived from the same muscle-type. Analysis details provided in Supporting Information.

### Myofibril Mechanics

Myofibrils from rat SOL and EDL muscles were prepared as described in the supplemental Material and Methods. In brief, myofibrils were attached between a rigid glass needle and a pre-calibrated atomic force cantilever that served as the force transducer within a microfluidic chamber in which calcium was varied between pCa 9.0 and 4.5 for tension:pCa measurements. The rate of force redevelopment following a large amplitude sarcomere length reduction (K_tr_) at pCa 4.5 was determined as a measure of actomyosin kinetics.

### Immunofluorescence

Immunofluorescence staining was performed on isolated myofibrils and cryo-sectioned muscle tissue. All buffers were diluted into phosphate buffered saline (PBS) at 4°C unless stated otherwise. SOL and EDL muscles were sectioned at 10 µm thickness using a cryostat microtome. Cryostat sections were incubated with 0.1M glycine (Sigma) followed by a 10% goat serum-based blocking buffer immediately prior to antibody labeling. Samples were incubated with primary antibodies against either slow (MYBPC1) or fast (MYBPC2) MyBP-C, and α-actinin (Sigma) for 2 hours. The sections were then rinsed with PBS and incubated with species specific Alexa Fluor secondary antibody for 2 hours. After the secondary incubation step, the sections were rinsed with PBS, mounted in a fluorescence anti-fading medium and sealed with nail polish. Imaging was performed on a Nikon Ti Eclipse inverted microscope equipped with a LU-N4/N4S 4-laser unit (405 nm, 488 nm, 561 nm and 640 nm). The same protocol was used for myofibril staining. Myofibril isolation was performed as previously described in (49). For more detail see supplemental Material and Methods.

### Immunofluorescence Image Analysis and Modeling

Briefly, we decomposed confocal immunofluorescence images into a set of one-pixel-wide intensity line scans then aligned these to each other by maximizing the pairwise cross-correlation. The aligned intensity line scans were fitted with a double Gaussian to determine the peak-to-peak spacing of the fluorescence doublets. We then developed an analytical model to predict the molecular arrangements that resulted in the dual Gaussian fluorescence profiles. The model accounted for the binding of MyBP-C at specific sites along each half of the thick filament, as described in the Results, and the point spread function of the fluorophores. We compared the model results to the data using Kolmogorov–Smirnov test, where p value >0.01 demonstrates significant overlap, to maximize the goodness-of-fit. For more detail see supplemental Material and Methods.

### Monte Carlo Simulations of Native Thick Filament Motility

Briefly, a Monte Carlo style simulation of thin filament motility over native thick filaments was used to generate 1000 displacement-vs-time trajectories for each set of possible model parameter values. Velocity distributions resulting from each of these simulations were then compared to experimental results to determine parameters that offer best agreement between simulation and experiment. For details see Supplemental Information.

## Footnotes

### Author contributions

A.L., S.N. M.J.P. and D.M.W. designed research; A.L., S.N, S.R. A.S.C., T.S.O., F.B. and S.B.P. performed research; S.S. and J.W.M. contributed new reagents/analytic tools; A.L., S.N., S.R., A.S.C., T.S.O., D.E.R, and M.J.P analyzed data. A.L., S.N., M.J.P and D.M.W wrote the paper with contributions from F.B, S.R., and T.S.O.

## Acknowledgements

The authors are indebted to Ms. Nicole Bouffard (Microscopy Imaging Center) and Mr. Todd Clason (Imaging / Physiology Core Facility) of the University of Vermont (USA) for their expert assistance in cryo-section and confocal imaging. The authors are also grateful to Prof. Doug Taatjes of the University of Vermont (USA) for his proficient advice on immunolabeling. This work was supported by National Institutes of Health (NIH) grants to D.M.W. (HL059408, AR067279 and HL126909), S.S. (HL130356, HL139680, AR067279, and HL105826), and M.J.P. (HL124041); American Heart Association funds to S.S. (19TPA34830084 and 19UFEL34380251) and J.W.M. (17POST33630095); and by funds from the Natural Sciences and Engineering Research Council of Canada to D.E.R, who is a Canada Research Chair (Tier I) in Muscle Biophysics.

