## Supplemental Information for "SKELETAL MyBP-C ISOFORMS TUNE THE MOLECULAR CONTRACTILITY OF DIVERGENT SKELETAL MUSCLE SYSTEMS"

**Supporting Information (SI Appendix)**
**MATERIALS & METHODS**

### SKELETAL MYBP-C ISOFORMS TUNE THE MOLECULAR CONTRACTILITY OF DIVERGENT SKELETAL MUSCLE SYSTEMS

Amy Li[1], Shane Nelson[1], Sheema Rahmanseresht[1], Filip Braet[2,3], Anabelle S. Cornachione[4], Samantha Beck Previs[1], Thomas S. O'Leary[1], James W. McNamara[6], Dilson E. Rassier[5], Sakthivel Sadayappan[6], Michael J. Previs[1], David M. Warshaw[1]

[1]Department of Molecular Physiology and Biophysics, University of Vermont, Burlington, VT 05405 USA.
[2]Discipline of Anatomy & Histology, School of Medical Sciences, The University of Sydney, Sydney, NSW 2006 Australia.
[3]Australian Centre for Microscopy & Microanalysis, The University of Sydney, Sydney, NSW 2006 Australia.
[4]Department of Physiological Science, Federal University of São Carlos, São Carlos, Brazil.
[5]Department of Kinesiology and Physical Education, McGill University, Montreal, Canada.
[6]Heart, Lung and Vascular Institute, Department of Internal Medicine, Division of Cardiovascular Health and Disease, University of Cincinnati, Cincinnati, OH 45221 USA

#### ANIMAL HANDLING AND ETHICS

All protocols were approved by the Institutional Animal Care and Use Committee of the University of Vermont and McGill University, and were in compliance with the Guide for the Use and Care of Laboratory Animals published by the National Institutes of Health. Sprague Dawley rats, 12-14 weeks of age and weighing 300 ± 40 g (Charles River Laboratories International, Quebec, CA), were kept under pathogen-free conditions and had free access to standard food and water. The animals were allowed to acclimatize for at least 72 h prior to use at the animal care facility prior to being euthanized using 3% isoflurane (50 mg/mL; NDC 57319-507-06, Phoenix Pharmaceuticals Inc., USA). When no longer responsive to a hard pinch to the feet, rats were decapitated with a small animal guillotine. The skin of the lower limbs was immediately resected to reveal the underlying muscle structures. The slow-twitch soleus (SOL) and fast-twitch extensor digitorium longus (EDL) muscles were identified and dissected.

#### PROTEIN ISOLATION FROM SKELETAL MUSCLE TISSUE

Myosin, native thick filaments and native thin filaments (NTF) were isolated from the rat SOL or EDL muscles using methods previously employed by our laboratory (1, 2). Unless otherwise indicated, experiments were performed using proteins isolated from the same muscle-type. Actin was prepared from chicken pectoralis (3). Actin and NTFs were fluorescently labeled with rhodamine phalloidin (4). All protein isolation procedures were performed at 4$^oC$ unless stated otherwise.

##### Myosin

Whole myosin was freshly isolated using methods previously employed by our laboratory (5). ~30 mg skeletal muscle tissue was homogenized in myosin extraction buffer (0.3 M KCl, 0.15 M $K_2HPO_4$, 0.01 M $Na_4PO_7$, 0.001 M $MgCl_2$, 0.001 M ATP and 0.002 M DTT, pH 6.8) for 15 min. The homogenate was allowed to sit at 4$^oC$ for 1 h with constant stirring to maximize yield. This was followed by ultracentrifugation for 1 h at 150,000 x g (TLA-120.1 rotor, Beckman Coulter). Filamentous myosin was precipitated from the supernatant and pelleted by ultracentrifugation for 20 min at 50,000 x g (MLS-50 rotor, Beckman Coulter) and resuspended in myosin storage buffer (0.6 M KCl, 0.002 M DTT, 0.02 M imidazole at pH 6.8, and in 50% glycerol). Before each experiment, the isolated myosin was ultra-centrifuged at 95,000 x g in the presence of actin and ATP to remove inactive myosin molecules from the preparation. Following this procedure, the myosin concentration was determined by a Bradford assay (6) using commercially available reagents from Bio-Rad Laboratories.

##### Native thick filament isolation

For native filament isolation, SOL or EDL muscle bundles were washed in relaxing solution (50mM NaCl, 5mM MgCl2, 2mM EGTA, 1mM DTT, 7mM phosphate buffer at pH 7, 10mM creatine phosphate, 5mM ATP) (2). The muscle was initially teased into 2-3 mm bundles under relaxing solution, and then into single muscle fibers with three fresh changes of skinning solution (relaxing solution containing 0.5% Triton-X 100).

Native thick filaments were enzymatically digested from muscle fibers with calpain-1 from porcine erythrocytes (CALBIOCHEM). 4-6 skinned fibers were minced in either 0.05 or 0.3 U/µL of calpain-1 in a buffer of 20mM imidazole (pH 6.8), 5µM 2-mercaptoethanol, and 1mM calcium acetate on a slide previously coated with 1 mg/mL of bovine serum albumin. The released native thick filaments were collected by a series of rinses with relaxing solution containing 75 mM NaCl. Native thick filaments relieved under conditions containing 0.3 U/µL calpain-1 selectively cleaved the N-termini off slow MyBP-C and was therefore used as control to determine the effects of the three slow MyBP-C isoforms.

**Native thin filament isolation**

For NTFs, all ultracentrifugation steps were undertaken using the TLA-100 rotor (Beckman Coulter). SOL or EDL muscle bundles underwent the same initial relaxing and skinning step as described in the native thick filament isolation. Immediately after skinning, the muscle fibers were homogenized in homogenization buffer (0.1 M KCl, 0.005 M MgCl2, 0.001 M EGTA, 0.025 M imidazole at pH 6.45, 0.01 M DTT, 0.005 M ATP, and 0.1% Triton X-100) for 15 min (1, 7). The homogenate was clarified by ultracentrifugation at 40,000 × g for 20 min. The supernatant was then centrifuged at 200,000 × g for 45 min. The pellet was resuspended in buffer containing 0.1 M KCl, 0.005 M MgCl2, 1 mM EGTA, 0.025 M imidazole (pH 7.9), 0.01 M DTT, and 0.005 M ATP; followed by centrifugation for 5 min at 40,000 × g. The supernatant was pelleted by ultracentrifugation for 45 min at 200,000 × g. The resulting pellet contained NTFs that was allowed to soften overnight with homogenization buffer without ATP or Triton X-100. The protein concentration of NTFs was determined using a Bradford assay, and was then fluorescently labeled at equimolar ratio with tetramethyl-rhodamine-phalloidin. The labeled NTFs were diluted to 5 nM in actin buffer (AB) containing 0.025 M KCl, 0.001 M EGTA, 0.010 M DTT, 0.025 M imidazole, 0.004 M $MgCl_2$, (pH 7.4), 70 µg/mL glucose oxidase, 45 µg/mL catalase, 5.834 mg/mL glucose.

**QUANITATIVE MASS SPECTROMETRY**

**Thick filament sample preparation for mass spectrometry**

Thick filaments samples were collected in 1.5 ml microcentrifuge tubes in 75 µM relaxing buffer and digested in-solution or proteins separated on 12% SDS Bis-Tris Protein gels (NuPAGE).

For in-solution digestion, a 75 µL aliquot of 0.1% RapiGest SF Surfactant (Waters Corporation) was added to each tube and the tubes heated at 50 °C for 1 hour. Proteins were reduced by addition of 5 µL of 0.1 M dithiothreitol to each tube and heating (100 °C, 10 min). Cysteines were alkylated by addition of 10.4 µL of 100 mM iodoacetamide in 50 mM ammonium bicarbonate (22±1 °C, 30 min, in dark). Peptides were generated by addition of 25 µL of 50 mM ammonium bicarbonate containing 5 µg of trypsin (Promega) (37 °C, 18 hours). The samples were dried down in speed vacuum device. The trypsin was deactivated and RapiGest cleaved by addition of 100 µL of 7% formic acid in 50 mM ammonium bicarbonate (37 °C, 1 hour). The resultant peptides were dried down and RapiGest was cleaved again by addition of 100 µL of 0.1% trifluoroacetic acid (37°C, 1 hour). The resultant peptides were dried down and reconstituted a final time in 150 µL of 0.1% trifluoroacetic acid. The tubes were centrifuged at 14,000 rpms for 5 minutes to pellet the surfactant and the top 125 µL of solution was transferred into a mass spectrometry (MS) analysis vial.

For in-gel digestion, large areas of the gels (150-75kDa) were excised from the gels, minced, destained in 50% acetonitrile and dehydrated using 100% acetonitrile and drying in a SpeedVac concentrator. Proteins were reduced by addition of 100 µL of 0.01 M dithiothreitol to each tube and heating (50 °C, 45 min). Excess dithiothreitol was removed and cysteines were alkylated by addition of 100 µL of 55 mM iodoacetamide in 50 mM ammonium bicarbonate (22±1 °C, 30 min, in dark). The solution was removed and the samples were washed with 3, 400 µL exchanges of 50% acetonitrile, with 15 minutes between exchanges. The samples were dried in the SpeedVac. Peptides were generated by addition of 100 µL of 50 mM ammonium bicarbonate containing 2 µg of trypsin (Promega) (37 °C, 18 hours). The samples were dried down in speed vacuum device and 100 µL of 7% formic acid in 50 mM ammonium bicarbonate was added to deactivate the trypsin. The resultant peptides were extracted with 3, 100 µL aliquot of 50% acetonitrile, with 15 minutes between exchanges. The aliquots were collected, dried down, reconstituted in 30 µL of 0.1% trifluoroacetic acid, and transferred into a mass spectrometry (MS) analysis vial.

##### Liquid chromatography mass spectrometry

A 20 µL aliquot of each sample was injected onto an Acquity UPLC HSS T3 column (100 Å, 1.8 µm, 1 × 150 mm) (Waters Corporation) attached to a UltiMate 3000 ultra-high pressure liquid chromatography (UHPLC) system (Dionex). The UHPLC effluent was infused into a Q Exactive Hybrid Quadrupole-Orbitrap mass spectrometer through an electrospray ionization source (Thermo Fisher Scientific). Data were collected in data-dependent MS/MS mode with the top five most abundant ions being selected for fragmentation.

##### Mass spectrometry data analyses

Peptides were identified from the resultant MS/MS spectra by searching against a rat proteome database downloaded from UniProt and a custom database using SEQUEST in the Proteome Discoverer 2.2 (PD 2.2) software package (Thermo Fisher Scientific). Slow-type MYBPC1 and fast-type MYBPC2 sequences were downloaded from the NCBI protein database (https://www.ncbi.nlm.nih.gov/protein). For slow-type MYBPC1, 16 unique rat sequences were identified, with differences consistent with alternative splicing of exons 3, 4, 5, and 30-33. In humans, additional splicing in exons 1-6, 10 and 23 has been identified. Therefore, sequences for hypothetical rat splice variants were created from the full-length slow-type MYBPC1 sequence, using all possible splicing patterns. The potential loss of methionine from the N-terminus of each protein (-131.20 Da), the loss of methionine with addition of acetylation (-89.16 Da) and addition of carbamidomthyl (C; 57.02 Da), oxidation (M, P; 15.99 Da: M; 32.00 Da) were accounted for by variable mass changes. Areas under each LC peak were calculated using PinPoint Software (Thermo Fisher Scientific). Peptide abundances were exported to Excel (Microsoft) and normalized by dividing each raw abundance by the mean abundance determined from 3 peptides shared between skeletal myosin isoforms, being LDEAEQIALK, DIDDLELTLAK and LASADIETYLLEK. Normalization corrects for differences for the amount of sample digested and injected onto the UHPLC column as previous described (8). The normalized

LC peak abundances were used of individual peptides of interest were used to determine relative protein ratios by a mass-balance label-free approach (8).

#### EXPRESSION OF N-TERMINAL FRAGMENTS

Recombinant proteins of the skeletal MyBP-C isoforms identified by LCMS were expressed and purified using a pET28 expression system (Millipore 70777) in E. coli with an N-terminal His-tag as previously described in (1). Briefly, the N-terminus of fast-type MyBP-C (fC1C2) and slow-type MyBP-C (sC1C2, sC1C2$_{\Delta exon4\text{-}5}$ and sC1C2$_{\Delta exon1\text{-}6.5}$) generated up to the C2 domain to be examined in our conventional native thin filament motility assays. We attempted to purify sC1C2$_{\Delta exon1\text{-}6.5}$ from both E. coli and Bacculovirus-Sf9 expression systems, however, this protein did not remain soluble without the presence of guanidine hydrochloride.

#### CONVENTIONAL NATIVE THIN FILAMENT MOTILITY ASSAY

For conventional *in vitro* motility assays, the motion of NTFs over myosin incubated on nitrocellulose-coated flow cell were observed by epifluorescence microscopy as described in (1). Briefly, skeletal myosin from rat SOL or EDL (100 µg/mL) was incubated in the flow cell (2 minutes), followed by blocking with 1 mg/mL BSA in actin buffer. To remove non-functional myosins, the flow cells were incubated with 1 µM unlabeled NTFs in AB for 1 minute, and then washed twice with 1 mM ATP in AB, followed by two washes with AB. Labeled NTFs were added to the flow cell for 1 minute and rinsed with AB. NTF motion was initiated by the addition of motility buffer comprised of AB with 100 µM ATP, 0.5% methyl cellulose, and $CaCl_2$. The motility buffer contained a range (pCa 4-9) of calcium concentrations as determined using MaxChelator software (9).

Motility experiments were performed using a Lumen 200W metal arc lamp (Prior Scientific), Nikon Eclipse TiU microscope with Plan Apo objective (100x, 1.35 n.a.) and Mega Z 10 bit digital camera (Stanford Photonics) for fluorescent excitation of labeled NTFs and image acquisition, respectively. Images were acquired using the software, Piper, at 10 frame/s without pixel binning (95 nm/pixel) and then down-sampled to 2 frames/s using ImageJ 1.51r (NIH).

The velocity of individual NTFs and percentage of mobile NTFs in each movie were analysed using DiaTrack 3.03 software (Semasopht). Data for thin filament velocity and the percentage of moving filaments were obtained. These two parameters were multiplied to give an effective velocity that takes into account the activation state of the NTF. This effective velocity value was then normalized to control. Data are presented as the mean normalized regulated velocity ± SEM from triplicate independent experiments.

Initial pCa-velocity experiments used to determine the optimal pCa to use for subsequent $Ca^{2+}$-dependent motility studies were plotted as velocity versus pCa and fitted to a sigmoidal dose-response curve (Fig. S4A), with pCa50 representing changes in calcium sensitivity. Two pCa concentrations were selected to represent NTFs that were effectively "off" (i.e., pCa7.5) and maximally "on" (i.e., pCa5). Subsequent experiments examined NTF motion at pCa5 and pCa7.5 as a function of increasing concentrations MyBP-C (Figs. 5, S6). These data were plotted as normalized regulated velocity against MyBP-C fragment concentrations (0-2 µM) (Fig. 5). All experiments were performed at room temperature. All data are presented as means ± SEM.

#### NATIVE THICK FILAMENT MOTILITY ANALYSIS

We developed a novel approach for the unbiased tracking of all fluorescent spots (i.e. shredded NTFs) in the thick filament motility assay.

1) All fluorescent spots were identified using the ThunderSTORM plugin (10), filtering for spots with integrated intensity > 2500 and a sigma value < 400nm. This excludes elongated and out-of-focus spots. These criteria identified approximately 100 fluorescent spots per frame.
2) Individual spot localizations were collected into Trajectories by linking spots that persist (to within 200nm frame-to-frame displacement) for a minimum of 10 consecutive frames.
3) An NTF shard is expected to travel in a straight line for approximately 800nm along the length of a half thick filament, thus, only trajectories meeting the following criteria were retained for further analysis:
    a. Aspect ratio of XY coordinates of > 2.0, as determined by Principal Component Analysis

b. 700nm < total end-to-end length < 900nm

4) Displacement (D) for each frame (i) from the first frame ($x_1$, $y_1$) was calculated as the Pythagorean distance from the first frame:

$$D_i = \sqrt{(x_i - x_1)^2 + (y_i - y_1)^2}$$

5) To detect a change in velocity, Displacement vs Time plots were then fit with 2 straight line segments, which we term a "dual-phase velocity". This was done using the segmented model regression function, implemented in the "Segmented" library for statistical programming language R
6) Trajectories were regarded as a single-phase if the dual-phase fit failed to satisfy the following criteria:
   a. Dual-phase fit must converge to a unique solution (as defined by the regression algorithm)
   b. Lifetimes of both phases must be > 5 frames (167ms)
   c. Difference between velocity of the two phases > 10%
7) If the velocity of either phase is zero ($p > .05$), that segment is discarded and the velocity of the remaining segment is retained as a "single phase" trajectory.
8) Finally, to exclude trajectories with highly variable velocities and/or poor tracking, we also calculated a "smoothness parameter," which we define as the coefficient of variance (standard deviation divided by mean) for the instantaneous velocity, as calculated over a 100ms sliding window. Trajectories (single- or double-phase) were excluded from analysis if their smoothness parameter > 2.0.

#### IMMUNOFLUORESCENCE

##### Tissue Processing

Whole SOL and EDL muscles were carefully dissected and briefly rinsed with phosphate-buffered saline (PBS) solution at pH 7.4 before placing the tissue in 4% paraformaldehyde in PBS at 4°C (i.e., fixative solution). Tissue samples were next processed for immunolabeling as described below.

#### Cryosectioning

Within 30 min of harvesting the tissue, the samples were transferred into a petri dish that contained ice-cold fixative solution. Next, 1 mm × 1 mm × ~8 mm tissue strips were cut—under fixative solution—along the long axis of the muscle using an ethanol-cleaned razor blade. The tissue strips were next washed three times for 3 min each with PBS at 4°C to remove excessive fixative, then excess buffer was removed using filter paper and transferred next into a plastic freezing mold filled with Optimum Cutting Temperature compound (Tissue Tek® O.C.T. Compound, Sakura, Cat No 4583, USA). Next, the freezing- and subsequent sectioning protocols detailed by Frederiks et al. (11) and Kumar et al. (12) were followed. Typically, tissue samples were embedded flat into the mold (i.e., normal functional orientation) and topped up with Tissue Tek® O.C.T. Compound. Immediately thereafter, samples were transferred individually to a dewar vessel containing pre-chilled iso-pentane (~160°C) cooled with liquid nitrogen. Briefly, the mold containing the embedded muscle tissue was held just above the cryogen for 10 sec before complete submersion into liquid iso-pentane for about 12 up to 15 sec. After freezing, 10 µm-thick cryo-sections at -18°C were cut using a cryostat microtome (HM 505N, MICROM International GmbH, Germany) equipped with a stainless steel knife (High Profile 818 Blades, Leica, Germany). Sections were collected and mounted on adhesive microscope slides (TruBond 380, Cat No #0380W, Matsunami, USA) and stored at -80°C in the dark upon immunostaining.

#### Immunolabelling

Cryostat sections were allowed to thaw to room temperature ($R_T$) for 15 min in a water humidified immunostaining chamber. The sections were next incubated in PBS buffer enriched with 0.1M glycine (Cat No G7126, Sigma-Aldrich, USA) at pH 7.4 for 10 min to quench free aldehyde groups, followed by a 40 min incubation with blocking buffer consisting of PBS, 10% normal goat serum (Cat No 005-000-121, Jackson Immunoresearch Laboratories Inc., USA), 0.1 M glycine and 0.1% Triton X100 (Cat No X100, Sigma-Aldrich, USA) at pH 7.4. Note that all above priming steps, including subsequent labeling steps, were executed at $R_T$ and within a fully humidified environment.

Sections were subsequently washed 5-times for 5 min each with PBS followed by a 2 hr incubation step with primary antibodies. Rabbit anti-MYBPC1 antibodies at 1:200 (Cat No 6679 [1 mg/mL], ProSci, USA) and mouse anti-MYBPC2 antibodies at 1:5 (Cat No MF1 [52 µg/mL], Hybridoma Bank, USA) were employed concurrently with anti-α-actinin antibodies for double-labeling purposes. Depending on the source of species the various primary anti-MYBPCs antibodies were raised, mouse anti-α-actinin (Cat No A7732 [1mg/mL], Sigma-Aldrich, USA) or rabbit anti-α-actinin (Cat No A2543 [71mg/mL], Sigma-Aldrich, USA) at a dilution of 1:500 in PBS was used: (i) to assess sample preservation, including overall labeling efficiency (13); and (ii) to define muscle fine structure using the Z-band staining pattern as a structural orientation marker (14). After the primary antibody incubation step, the samples were rinsed 5-times for 5 min each with PBS followed by a 2 hr incubation step with secondary antibodies. Taking the species origin of the primary antibodies into account, the fluorescent signal was attained using the corresponding Alexa Fluor™ secondary divalent antibody fragments (Invitrogen - Thermo Fisher Scientific, USA): i.e., goat anti-mouse 488 (Cat No A11017), goat anti-mouse 647 (Cat No A21237), goat anti-rabbit 488 (Cat No A11070) and goat anti-rabbit 647 (Cat No A21246). After secondary antibody labeling, the samples were rinsed 5-times for 5 min each with PBS and immediately mounted in anti-fading medium (Fluorescence Mounting Medium, Cat No S3023, Dako, USA). Slides were next covered with cover glasses (Micro Cover Glass 24 × 30 mm - No 1.5, Cat No 48404-466, VWR, USA), sealed with transparent nail polish and stored at 4°C in the dark upon confocal laser scanning microscopy (CLSM) examination (see, *vide infra*).

**Confocal Laser Scanning Microscopy**

Immunolabeled tissue sections were examined with a Nikon C2 CLSM installed on a Nikon Ti Eclipse inverted microscope using a Plan Apo 60× / 1.4 N.A. oil objective lens. The microscope system was equipped with a LU-N4/N4S 4-laser unit (405 nm, 488 nm, 561 nm and 640 nm), an external phase contrast unit, a fully automated motorized stage, and was running under the Nikon TCS-NT software (version 4.5). Confocal data sets were acquired typically at 1024 × 1024 pixels and/or 2048 × 2048 pixels at a scanning speed of 5.2 msec/l and 9.4 msec/l, respectively. The performance of the microscope was regularly assessed using TetraSpeck™ calibration-grade fluorescent microspheres mounted on a slide (Cat No T14792, Invitrogen - Thermo Fisher Scientific, USA).

**Image and Statistical Analysis**

To determine the MyBP-C doublet spacing in confocal images, we selected a region of interest (ROI) of approximately 20x10 microns for each labelling condition. Within each ROI, intensity linescans were spatially aligned using the following approach:

1) An initial estimate of average sarcomere length was determined as the maximum of the autocorrelation function, as calculated on a 1-pixel wide intensity linescan. This was usually near 2 microns in length.
2) Individual intensity profiles were collected as separate, 1-pixel wide intensity linescans, 1.5 sarcomeres in length, each centered near the middle of the MyBP-C doublet. This was done over all rows in the ROI, resulting in 300-400 separate intensity profiles.
3) Each of these repeats was then aligned to an aggregating "seed" profile by finding the spatial offset that maximizes the cross-correlation between each repeat and the seed. After alignment of each individual profile, the seed is (recursively) recalculated as the averaged profile of the aligned repeats. Sub-pixel refinement of the spatial offset value was accomplished by fitting a trigonal spline to a plot of cross-correlation vs spatial offset, with the local maximum interpolated from the fitted spline.
4) After spatial alignment of all repeats, the cloud of intensity vs aligned position was fit with a double Gaussian model:

$$I = \frac{A}{\sqrt{2\pi\sigma^2}} e^{-\frac{(x-m_1)^2}{2\sigma^2}} + \frac{A}{\sqrt{2\pi\sigma^2}} e^{-\frac{(x-m_2)^2}{2\sigma^2}}$$

Where $I$ is the Intensity, $m_1$ & $m_2$ is the center of the 2 MyBP-C densities and $\sigma$ is the width (Standard Deviation) of each peak. These fits were found to be superimposable with averaged (mean or median) profiles of the aggregated data.

**MODELING**

#### Analytical Modeling of MyBP-C Fluorescence

The fluorescence signals from immuno-fluorescently-labeled MyBP-C molecules in cryo-sectioned SOL and EDL muscles and quantitative mass spectrometry data were used in an analytical model to predict the localization and distribution of MyBP-C molecules within the average sarcomere. The model assumed that MyBP-C molecules can be located along each half of the thick filament in any of the 17 stripes associated with the myosin helical repeat that are spaced 43 nm apart. It also assumed a 200 nm bare zone between each half of the thick filament in a sarcomere. The models predictions included the number of MyBP-C stripes along the length of the thick filament, the location of these stripes, and the number of MyBP-C molecules within each stripe.

To generate theoretical images for the average fluorescence signal coming from multiple sarcomeres, each MyBP-C stripe was rendered with a 2-dimensional Gaussian distribution with a full-width-half-maximum (FWHM) of 400 nm. The intensity of Gaussian distribution represented the number of MyBP-C molecules in one stripe. The higher the intensity, the higher the number of MyBP-C molecules in that stripe. The model was constrained to include a total of 27 MyBP-C molecules per half of the thick filament, based on the total abundance of MyBP-C measured relative to myosin in the SOL and EDL muscle samples. We iteratively adjusted the number of stripes, the number of MyBP-C stripes along the length of the thick filament, the location of these stripes, and the number of MyBP-C molecules within each stripe, depending on the muscle type.

For each iteration, we generated a theoretical image and fitted the intensity profile along the length of the sarcomere with the sum of two Gaussian distributions, as was done for the average confocal images. The data from the average confocal images and fitted model were compared using Kolmogorov–Smirnov test. A p-value of > 0.01 assures the goodness-of-fit.

#### Monte Carlo Simulation

To reconcile the slowing of thin filament velocity by MyBP-C observed in both the conventional motility assay and the native thick filament assay, as well as an effort to infer the inhibitory influence of the insoluble, truncated slow-type MyBP-C$_{\Delta exon1\text{-}6.5}$ splice variant, we developed a Gillespie style Monte Carlo simulation of the SOL thick filament motility assay.

A half thick filament is defined by 17 myosin "stripes," each 43nm in length and numbered beginning with the one closest to the center of the thick filament (M-line). According to structural information (15, 16), slow-type MyBP-C is assigned to stripes 3-11, with the slow-type MyBP-C isoform in each stripe randomly assigned for each simulation, according to the experimentally observed relative abundances (Table 2). In the simulation, each stripe has an "Inherent Velocity" ($V_{inh}$) which we define as:

$$V_{inh} = \frac{V_{max}}{1 + \frac{I}{K_i}}$$

where the maximal velocity, $V_{max}$ is randomly drawn once for each trajectory from the experimentally observed 1st phase velocities (790±320nm/s, Table 3), and the $K_i$ is the inhibitory capacity of the specified slow-type MyBP-C isoform (Fig. 5, Table S1) , and $I$ is the effective local concentration of slow-type MyBP-C in this context (0 for stripes that do not contain MyBP-C).

Native thin filament shards are considered to be 250±50nm in length (2), and are tracked according to their center. The velocity of the NTF is specific to its position on the thick filament (see below). Coordinates along the thick filament are in nm, beginning with 0nm at the tip of the thick filament and 731nm (43nm/stripe x 17 stripes) for edge of stripe 1 nearest the M-line. Note that negative coordinates (where a portion of the NTF is in contact with the thick filament, but the center of the NTF is distal to the thick filament tip) are allowed in the simulation.

Each simulation run begins with a randomly generated starting position near the tip of the thick filament (drawn from a Gaussian distribution, mean=0nm, σ=20nm). Beginning with this initial position, subsequent NTF positions are calculated at 10nm intervals for locations ≤ 731nm. For each of these positions, the Effective Velocity ($V_{eff}$) is calculated as the mean of the Inherent Velocities ($V_{inh}$) of the thick filament stripes that are in contact with the NTF. From this Effective Velocity, a waiting time for each position is randomly generated from an exponential distribution with a rate of:

$$k_{wait} = V_{eff}/10 \text{ nm}.$$

A preliminary displacement vs time trace is then generated by plotting each position vs the cumulative waiting time for that position. Finally, this preliminary trace is then re-sampled to the experimental frame rate (30FPS) by binning the cumulative summed waiting times into non-overlapping 33ms bins and averaging the NTF position in a time-weighted manner within each bin. Tracking error, in the form of random values drawn from a Gaussian distribution (mean = 0nm, $\sigma$= 20nm) are added to the resampled position coordinates.

Within this simulation there are 2 free parameters: *I*, or the effective local concentration of sMyBP-C, relative to the concentration of recombinant sMyBP-C fragments added to the conventional motility experiments, and the $K_i$ for the truncated $sC1C2_{\Delta exon1\text{-}6.5}$ which was insoluble when expressed, precluding its testing in motility experiments. Therefore, we performed a systematic search over a range of plausible values for these two parameters. For each combination of values for these two parameters, two sets of simulations were run – one for each of the calpain concentrations that were used experimentally. This enzymatic treatment resulted in altered occupancy of the various MyBP-C variants (Table 2). In the simulation, stripes with MyBP-C that had been proteolytically digested by calpain are regarded as devoid of MyBP-C (*I* = 0mM). For a given condition, we simulated 1000 trajectories and then subjected those simulated trajectories to the same analysis used for experimental thick filament motility trajectories. The 6 resulting velocity distributions (Single phase, 1$^{st}$ & 2$^{nd}$ segments of double phase trajectories for both low- and high-Calpain treatment) were then compared to the experimental velocity distributions (Fig. 3, Table 3) using the “D” test statistic from the Kolmogorov-Smirnov test. Also, the fraction of single-phase runs was compared between the simulated and experimental results according to:

$$(\text{Simulation} - \text{Experiment})/\text{Experiment}.$$

This was done for both calpain treatment levels. Finally, an aggregate score for a given set of parameters was calculated as the sum of the 6 “D” statistics and the normalized fraction of single-phase runs for both calpain treatment levels. Lower aggregate scores (closer to zero) indicating closer agreement between simulated and experimental velocity distributions. As can be seen in Fig. 6, this simulation indicates a number of possible values that equivalently match the experimental results.

**MYOFIBRIL MECHANICS**

Small pieces of the EDL and SOL muscles were homogenized following standard procedures (17-19), which resulted in a solution containing isolated myofibrils. The myofibrils were transferred to a chamber placed on the top of an inverted microscope equipped with pre-calibrated atomic force cantilevers (AFC) used for force measurements (20). Myofibrils chosen based on striation pattern and number of sarcomeres in series (between 10 and 30) were attached between the AFC and a rigid glass needle. A computer-controlled, multichannel fluidic system connected to a double-barreled pipette was used for activation of the myofibril in solutions with different $Ca^{2+}$ concentrations, ranging from pCa 9.0 to pCa 4.5 (15°C). Length changes during the experiments were induced with a rigid micro-needle connected to a piezoelectric motor. Under high magnification, the contrast between the dark bands of myosin (A-bands) and the light bands of actin (I-bands) provided a dark-light intensity pattern, representing the striation pattern produced by the sarcomeres, which allowed measurements of sarcomere length during the experiments.

Once the myofibrils were fully activated and maximal force for a given $Ca^{2+}$ concentration was obtained, we induced shortening of the myofibrils (amplitude: 30% of sarcomere length; speed: 10 μm/s) during which the force declined and then rapidly redeveloped to reach a new steady state. The maximal force produced by the myofibrils was calculated after force development stabilization. Forces were averaged for a period of 2 s once steady state was reached. All forces were normalized by the myofibril cross-sectional area. For each contraction, the rate of force redevelopment ($K_{tr}$) following shortening was analyzed with a two-exponential equation (a*exp(-K*(t-c))+b), where t is time, K is the rate constant for force development, a is the amplitude of the exponential, and b and c are constants.

**Supporting Information (SI Appendix)**
**SUPPLEMENTARY FIGURES & TABLE**

### SKELETAL MYBP-C ISOFORMS TUNE THE MOLECULAR CONTRACTILITY OF DIVERGENT SKELETAL MUSCLE SYSTEMS

Amy Li[1], Shane Nelson[1], Sheema Rahmanseresht[1], Filip Braet[2,3], Anabelle S. Cornachione[4], Samantha Beck Previs[1], Thomas S. O'Leary[1], James W. McNamara[6], Dilson E. Rassier[5], Sakthivel Sadayappan[6], Michael J. Previs[1], David M. Warshaw[1]

[1]Department of Molecular Physiology and Biophysics, University of Vermont, Burlington, VT 05405 USA.
[2]Discipline of Anatomy & Histology, School of Medical Sciences, The University of Sydney, Sydney, NSW 2006 Australia.
[3]Australian Centre for Microscopy & Microanalysis, The University of Sydney, Sydney, NSW 2006 Australia.
[4]Department of Physiological Science, Federal University of São Carlos, São Carlos, Brazil.
[5]Department of Kinesiology and Physical Education, McGill University, Montreal, Canada.
[6]Heart, Lung and Vascular Institute, Department of Internal Medicine, Division of Cardiovascular Health and Disease, University of Cincinnati, Cincinnati, OH 45221 USA

**Supplementary Figure 1**. Slow- and fast-type MyBP-C peptides as identified by LCMS. The table shows the peptide sequence, the exons covered by the peptide, the domain in which the peptide is located and the isoform which the peptide is specific to. The alphabetical lettering on the extreme left-hand side of the table relates the schematic above the table, which links the peptide sequence to its location within each isoform. Peptides exist that are unique to a specific isoform, while other peptides are common amongst some or all isoforms. The arrowheads show the start and end of the peptide sequence on the exon map in Figs. S2 and S3.

**Supplementary Figure 2**. Exon mapping of slow- and fast-type MyBP-C N-terminal amino acids up to C2 domain. **(A)** The N-terminal amino acid sequence of the three slow-type MyBP-C were aligned, and the exons are mapped accordingly along the top. Green text denotes amino acids that are alternately spliced. Blue text denotes Ig domains. Red text denotes calpain cleavage site. Arrowheads denote start and end of the LCMS peptides used to detect isoform abundances and are color-coded in accordance with Fig S1. Accession numbers: sC1C2: NP_001094228.2, sC1C2$_{\Delta exon4\text{-}5}$ : XP_008763480.1, sC1C2$_{\Delta exon1\text{-}6.5}$ is a hypothetical sequence based upon the two preceding rat sequences, but beginning with the conserved alternate start codon utilized in the human sequence EAW97667.1 **(B)** Fast-type MyBP-C exons were mapped separately. Blue text denotes Ig domains. Accession number NP_001099727.1.

**Supplementary Figure 3**. Exon mapping of slow- and fast-type MyBP-C C-terminal amino acids. (**A**) Two slow-type MyBP-C C-terminal splice variants are presented. The C10 domain is denoted by blue text, and first and last amino acids of peptide sequences detected by LCMS (see Fig. S1) are indicated by the arrows. (**B**) Fast-type MyBP-C C terminus.

**Supplementary Figure 4**. Calcium-dependent *in vitro* motility assay and myofibril mechanics controls. (**A-F**) Slow-twitch SOL (grey) and fast-twitch EDL (black) native thin filaments (NTFs) were observed moving over their respective monomeric muscle myosins. (**A, D**) Velocity of NTF motion at varying calcium concentrations (pCa). (**B, E**) The NTF fraction moving (%) at each pCa. (**C, F**) The velocity x fraction moving. A-F are plotted for control (top panels) and sonication-shredded (bottom panels) NTFs. SOL and EDL NTFs exhibited similar sigmoidal responses to calcium (SOL $pCa_{50}$ = 6.59 ± 0.08, and EDL $pCa_{50}$ = 6.71 ± 0.02). The maximal sliding velocity of the SOL ($V_{max}$=0.48±0.03 µm/sec) NTFs is approximately half that of the EDL ($V_{max}$= 1.18 ± 0.04 µm/sec). Sonication was used to mechanically shred NTFs to ensure their lengths were sufficiently short (250±9nm, (2)) for accurate probing of the native thick filament regions with and without MyBP-C, C-zone and D-zone, respectively. Upon sonication, SOL and EDL NTFs remain fully regulated with calcium sensitivity (SOL $pCa_{50}$ = 6.69 ± 0.15, and EDL $pCa_{50}$= 6.59 ± 0.10) and maximal sliding velocities of (SOL $V_{max}$ = 0.44 ± 0.05 µm/sec, and EDL $V_{max}$ = 1.06 ± 0.07 µm/sec). All data are presented as mean ± SEM n=3 entire pCa curves per condition. (**G**) Tension normalized to myofibril cross-sectional area (T) vs. pCa for SOL and EDL myofibrils. Tension:pCa relationships were fitted with a Hill equation as follows: SOL: $T_{max}$=68.0±1.7 nN/µm$^2$, $K_m$=pCa5.9±0.02, $n$=2.4±0.3; EDL: $T_{max}$=58.2±3.0 nN/µm$^2$, $K_m$=pCa5.7±0.04, $n$=2.7±0.6 (parameter ± error of the fit). All data are presented as mean ± SEM, n=13 (SOL) and n=12 (EDL). (**H**) The rate of tension redevelopment ($K_{tr}$) following a rapid shortening of myofibrillar length at pCa 4.5 for SOL and EDL myofibrils. The faster $K_{tr}$ for EDL compared to the SOL myofibrils is reflected in the faster NTF velocities at pCa 5 in the motility assay (Figs. S4A,D). All data are presented as mean ± SEM, n=13 (SOL) and n=12 (EDL).

**Supplementary Figure 5**. Bacterial expression of the N-terminal $sC1C2_{\Delta exon1\text{-}6.5}$ fragment. $sC1C2_{\Delta exon1\text{-}6.5}$ appeared in the insoluble fraction using both BL21 (left) and Rosetta PLysS (right) cell lines. Arrows indicate the predicted migration and molecular weight of

$sC1C2_{\Delta exon1\text{-}6.5}$. Image is of a total protein coomassie stained gel. Although the fragment could be expressed, it was insoluble under the experimental conditions of the motility assay.

**Supplementary Figure 6**. Effects of recombinant slow- and fast-type MyBP-C N-terminal fragments on velocity and fraction moving of SOL and EDL native thin filaments. pCa5 data are shown as open data points and fitted with solid lines using non-competitive inhibitory curves (Table S1). pCa 7.5 data are shown as closed data points and fitted to Hill activation curves (dashed lines) (Table S1). Each fragment has a differential effect on (**A**) SOL thin filament velocity, (**B**) SOL thin filament fraction moving, (**C**), EDL thin filament velocity and (**D**) EDL thin filament fraction moving. These effects were assessed as a function of fragment concentration. All data presented as Mean ± SEM, n=4-5 curves per condition. See Table S1 for fit parameters.

**Supplementary Table 1**. Native thin filament motility parameters of fit to the Hill activation (pCa7.5) and non-competitive inhibition (pCa5) curves.

| | | Activation (pCa 7.5) | | | Inhibition (pCa 5) | | |
|---|---|---|---|---|---|---|---|
| | | $K_a$ (μM) | Normalized $V_{max}$ | n | $K_i$ (μM) | Normalized $V_{max}$ | n |
| SOL | sC1C2 | 0.23 ± 0.01 | 0.26 ± 0.02 | 4 | 3.77 ± 1.30 | 1.05 ± 0.06 | 4 |
| | sC1C2$_{\Delta exon4\text{-}5}$ | 0.62 ± 0.09 | 0.85 ± 0.08 | 4 | 119.9 ± 643.80 | 1.07 ± 0.05 | 5 |
| EDL | sC1C2 | 1.73 ± 4.78 | 0.27 ± 0.25 | 4 | 2.72 ± 0.48 | 1.04 ± 0.03 | 4 |
| | sC1C2$_{\Delta exon4\text{-}5}$ | 0.53 ± 0.10 | 0.76 ± 0.09 | 5 | 2.82 ± 0.72 | 1.02 ± 0.05 | 5 |
| | fC1C2 | 2.17 ± 8.20 | 0.03 ± 0.01 | 5 | 2.51 ± 0.43 | 1.03 ± 0.04 | 5 |

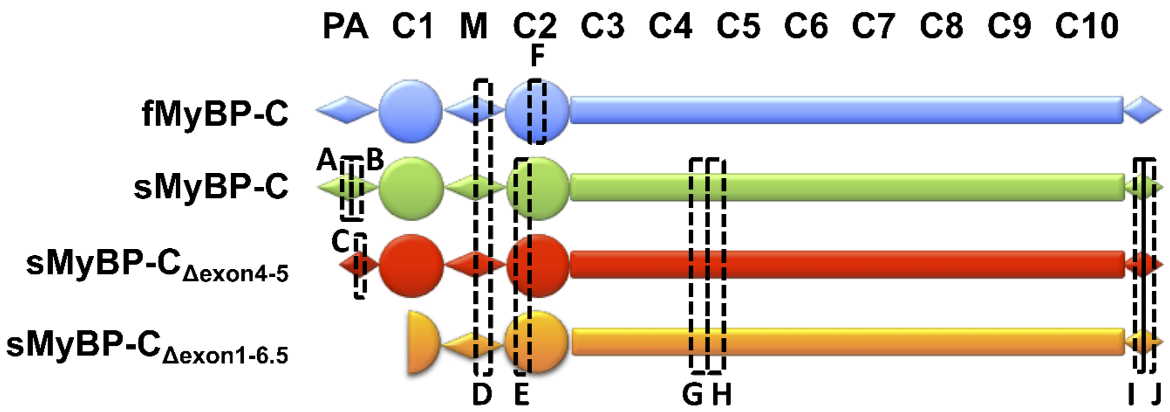


| | PEPTIDE | EXON | LOCATION | ISOFORM |
|---|---|---|---|---|
| A | EAGTAPAKEEDEASPPSALPPGLGSR | 3/4/5 | PA | sMyBP-C |
| B | DSEWSLGESPAGGEEQDKQNANSQLSTLFVE KPQTGSVK | 5/6 | PA | sMyBP-C |
| C | EWSLGESPAGGEEQDKQNANSQLSTLFVEKP QTGSVK | 3/6 | PA | sMyBP-C$_{\Delta exon4\text{-}5}$ |
| D | IAFQYGITDLR **(Common Peptide)** | 11 | M-DOMAIN | All MyBPC |
| E | ILDPAYQIDK | 12 | C2 | all sMyBP-C isoforms |
| F | LVVEISDPDLPLK | 9 | C2 | fMyBP-C |
| G | LSVDLRPLK | | C4 | all sMyBP-C isoforms |
| H | ITTPLTDQTVK | 16 | C4-linker-C5 | all sMyBP-C isoforms |
| - | EENEVPAPAPPPEEEDEASPPSALPPGLGSR | 1/2/4/5 | PA | sMyBP-C$_{\Delta exon3}$ |
| I | VIYQGVNTPGQPVSLEGQQQL | 31/32 | Post-C10 linker | sMyBP-C$_{\Delta exon33}$ |
| J | LLLQGVPPNIIDSYLR | 33 | Post-C10 linker | sMyBP-C$_{\Delta exon31\text{-}32}$ |
| J | GGLSFCR | 33 | Post-C10 linker | sMyBP-C$_{\Delta exon31\text{-}32}$ |
| - | LEVKGGLSFCRLLLQGVPPNIIDSYLR | 30/33 | Post-C10 linker | sMyBP-C$_{\Delta exon31\text{-}32}$ |
| J | DLQSSNPEEY | 33 | Post-C10 linker | sMyBP-C$_{\Delta exon31\text{-}32}$ |

**Supplementary Figure 1**. Slow- and fast-type MyBP-C peptides as identified by LCMS.

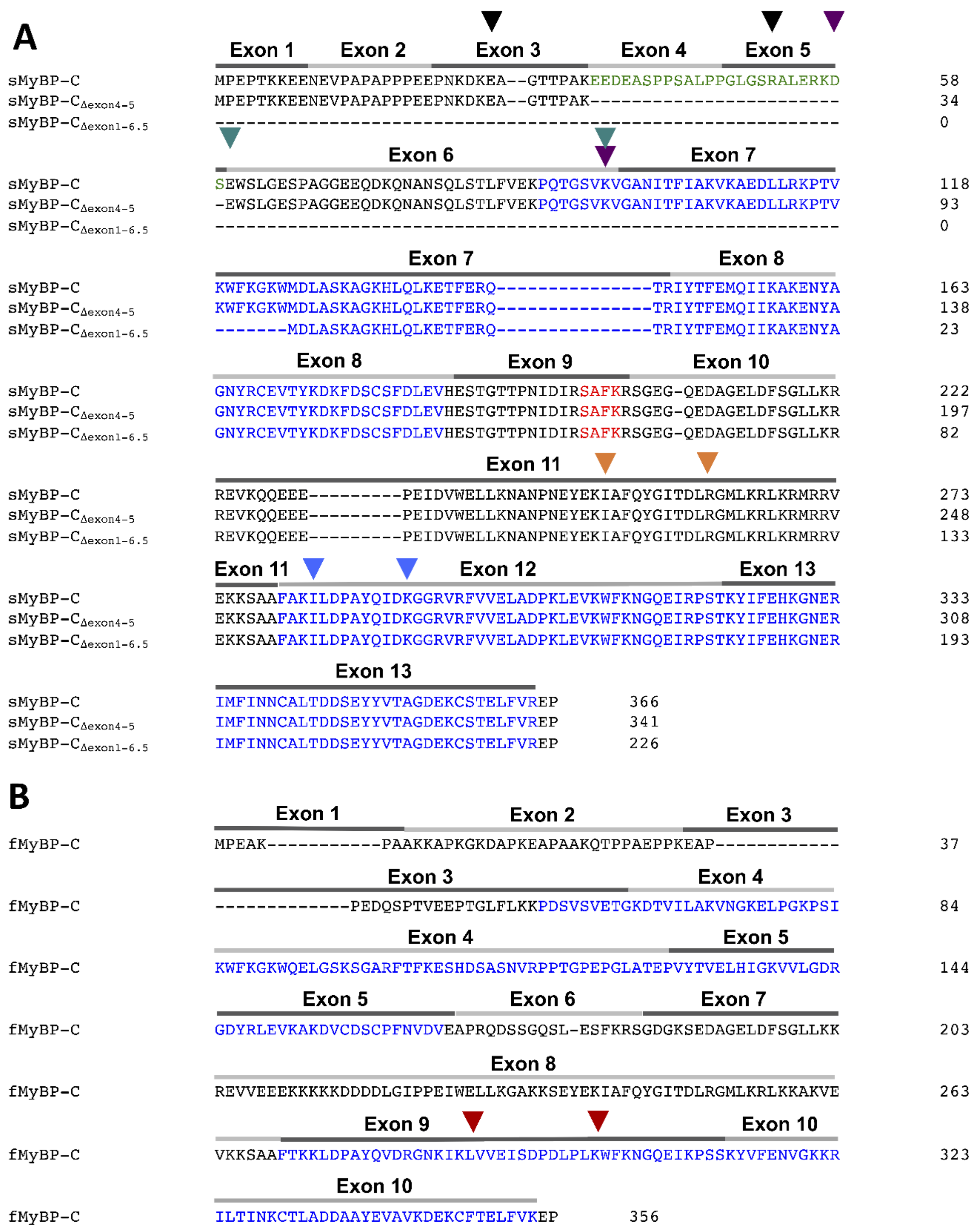

**Supplementary Figure 2**. Exon mapping of slow- and fast-type MyBP-C N-terminal amino acids up to C2 domain.

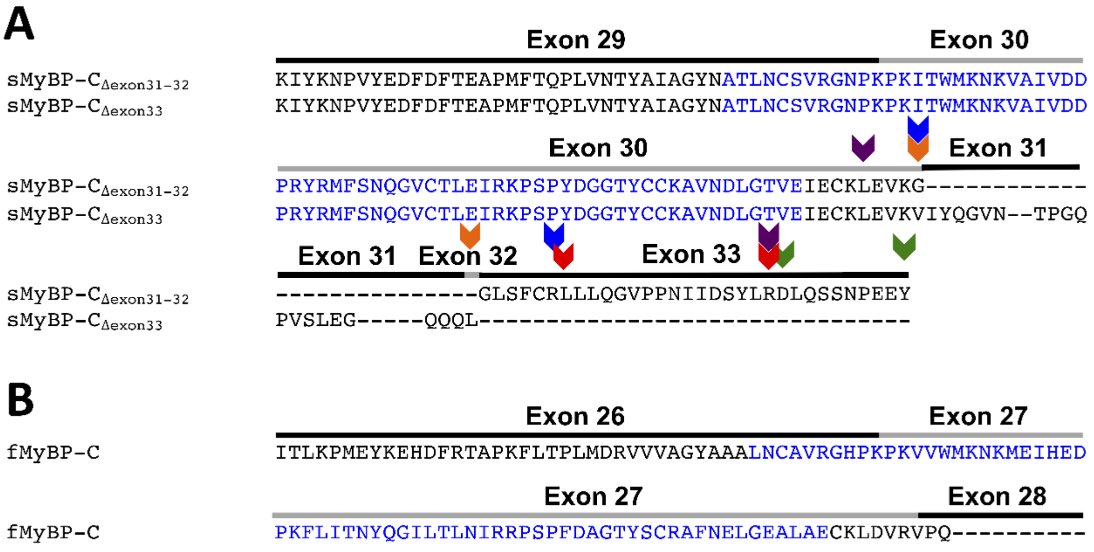


**Supplementary Figure 3**. Exon mapping of slow- and fast-type MyBP-C C-terminal amino acids.

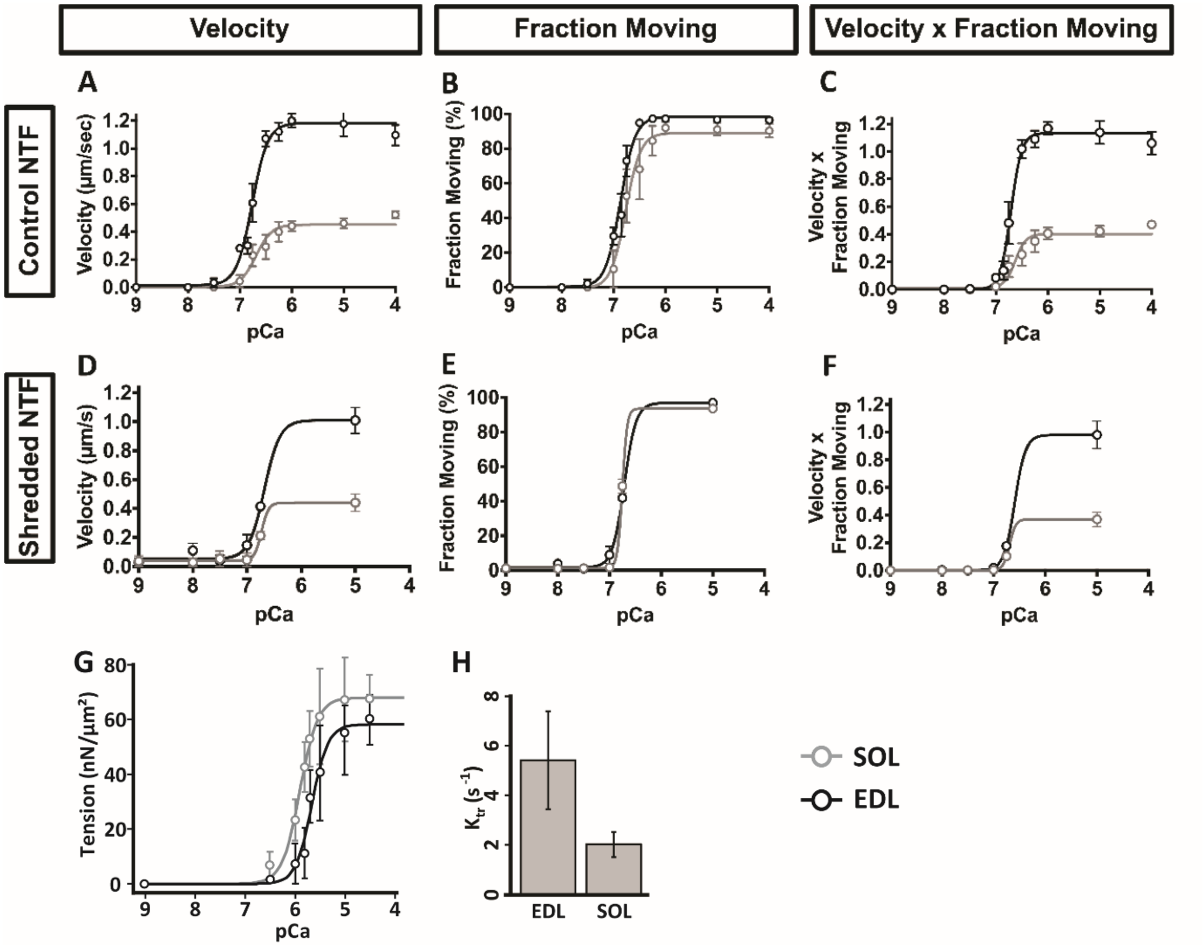


**Supplementary Figure 4**. Calcium-dependent *in vitro* motility assay and myofibril mechanics controls.

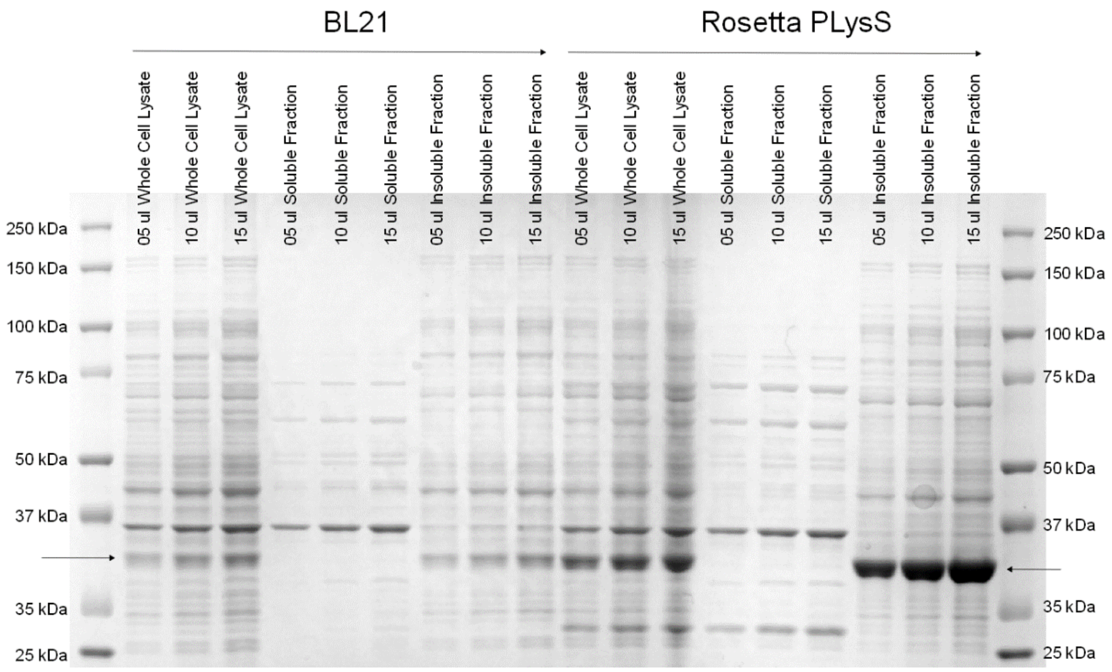


**Supplementary Figure 5**. Bacterial expression of sC1C2$_{\Delta exon1\text{-}6.5}$.

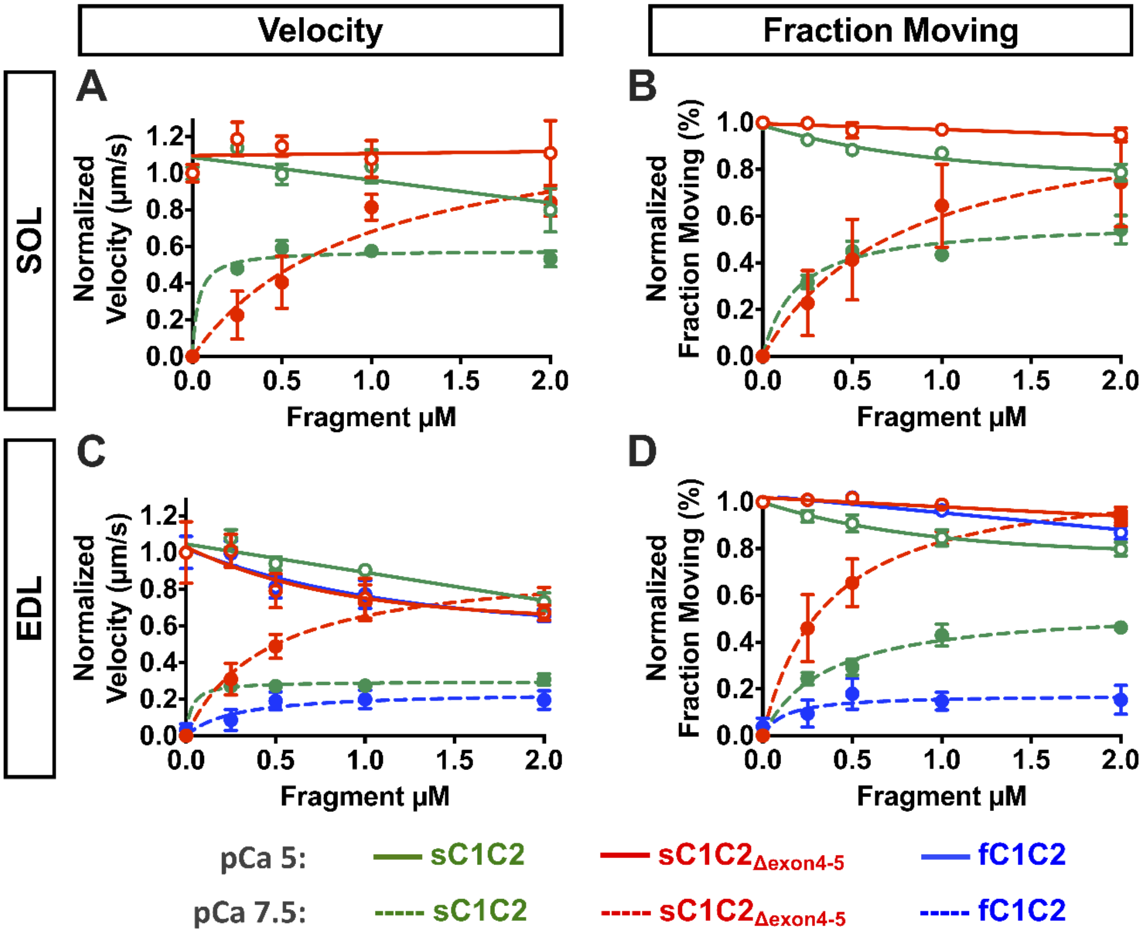


**Supplementary Figure 6**. Effects of recombinant slow- and fast-type MyBP-C N-terminal fragments on velocity and fraction moving of SOL and EDL thin filaments.
